# ExTrack characterizes transition kinetics and diffusion in noisy single-particle tracks

**DOI:** 10.1101/2022.07.13.499913

**Authors:** Francois Simon, Jean-Yves Tinevez, Sven van Teeffelen

## Abstract

Single-particle tracking microscopy is a powerful technique to investigate how proteins dynamically interact with their environment in live cells. However, the analysis of tracks is confounded by noisy molecule localization, short tracks, and rapid transitions between different motion states, notably between immobile and diffusive states. Here, we propose a probabilistic method termed ExTrack that uses the full spatio-temporal information of tracks to extract global model parameters, to calculate state probabilities at every time point, to reveal distributions of state durations, and to refine the positions of bound molecules. ExTrack works for a wide range of diffusion coefficients and transition rates, even if experimental data deviate from model assumptions. We demonstrate its capacity by applying it to slowly diffusing and rapidly transitioning bacterial envelope proteins. ExTrack greatly increases the regime of computationally analyzable noisy single-particle tracks. The ExTrack package is available in ImageJ and Python.

## Introduction

Studying the motion of proteins by single-particle tracking (SPT) allows to characterize how proteins dynamically interact with their environment (1, 2). Notably, single-particle tracking can reveal if and where proteins are diffusive or immobile (3–5). This information has significantly improved our understanding of important biological processes such as transcription-factor binding dynamics, antibody recognition, cytoskeletal dynamics, or intracellular transport (2, 6–13).

Molecules often transition between different motion states. If transitions happen rarely and if trajectories are long, different states such as immobile or diffusive states are reliably detected from time-averaged quantities such as the mean-squared displacement (MSD) (14–17). However, molecules often undergo rapid transitions between different states (5, 6, 10). Furthermore, tracks are often short as particles can bleach or diffuse out of the field of view or focal plane (17). In such situations, probabilistic methods are better suited to determine global parameters such as diffusion coefficients and transition rates (3, 7, 18–26). Some of these methods can also predict the motion states of individual molecules at every time point (3, 7, 26, 27), which can reveal the locations of binding sites, spatial correlations, and complex, potentially non-Markovian dynamics (28).

Previous probabilistic methods for diffusive models shown to correctly estimate diffusion and transition parameters (3, 25) are based on absolute distances between subsequent localizations. These methods have been developed for situations where physical displacements are large in comparison to the localization uncertainty for each molecule. However, when molecules transition rapidly between states, high time resolution is needed, which results in small physical displacements, which, in turn, make identifying different motion states hard or impossible (Fig. 1a-c). On the contrary, the whole track still allows the distinction of states (Fig. 1a), simply because subsequent positions of immobile or slowly diffusing molecules fall in the same small area determined by localization error, while subsequent positions of fast-diffusing molecules are nearly uncorrelated.

**Figure 1.**
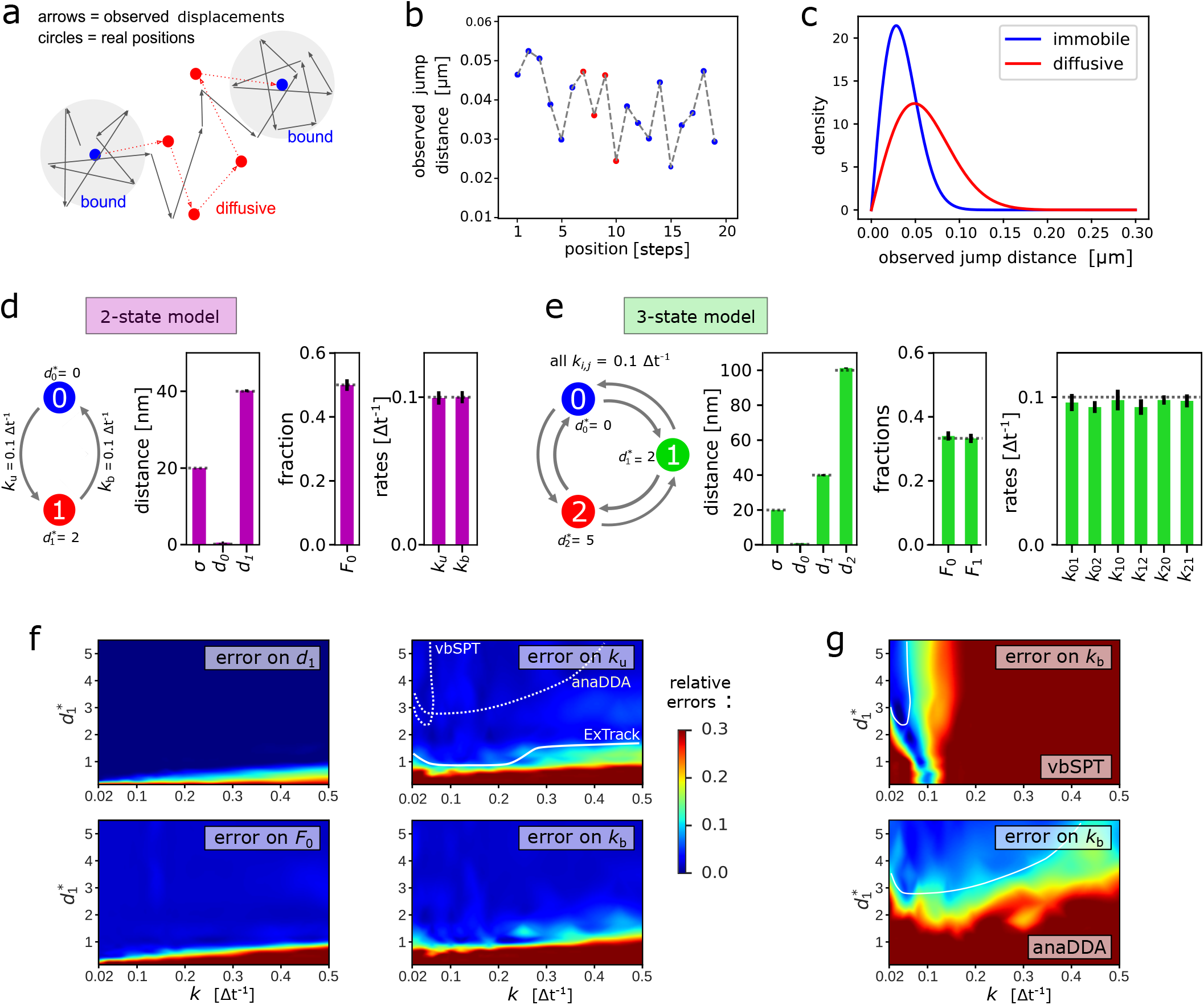
ExTrack permits to assess a wide range of multi-state diffusion models. **a**: Example track of a molecule transitioning between immobile and diffusive states with *d*_1_ = 2 *σ*. Arrows: observed displacements; dots: actual positions of immobile (blue) and diffusive (red) molecules. **b**: Consecutive observed distances of the track from **a. c**: Density function of observed distances of coefficiently immobile (blue) or diffusive (red) molecules for *d*_1_ = 2 *σ*. **d-e**: Left: Simulated two-state (**d**) and three-state (**e**) diffusion models with diffusion length and transition rates as indicated. Right: Model parameters estimated by ExTrack (mean *±* standard deviation) assuming a two-state (**d**) or three-state (**e**) model (localization error *σ*, diffusion lengths *d*_0_ and *d*_1_, initial immobile fraction *F*_0_, transition rates). Dotted lines: ground truth. ExTrack settings: two-state data: 2 sub-steps, window length = 10; three-state data: no sub-steps, window length = 7. **f**: Heatmap of the relative errors of *d*_1_, *F*_0_, *k*_*u*_ and *k*_*b*_ obtained from a two-state model fit to two-state simulations as in **d**. Error: mean absolute relative errors from 10 replicates per condition. White lines indicate regions of *<* 10% error for model parameters hardest to fit for ExTrack (*k*_u_, solid), vbSPT and anaDDA (*k*_b_, dashed, see **g**). **g**: Error on *k*_*b*_ of vbSPT and anaDDA (same protocol and color map as in **f**). See Fig S4 for errors on the other parameters.

To account for those spatial correlations, the full sequence of track positions must be taken into account. This approach has been used to characterize a single population of diffusing molecules (29, 30). However, if molecules transition between states, this approach becomes computationally demanding, because all possible sequences of singlemolecule states need to be considered. To avoid this computational complexity, different mean-field approximations (20, 21, 31–33) and machine learning approaches (34) have been proposed. However, their performance across model parameters remains to be investigated.

Here, we propose an alternative probabilistic method to extract diffusive motion states and transitions: We tackle the combinatorial problem of different motion states by introducing a sliding window that maintains the most important spatio-temporal correlations. The method is fast and accurate for a large range of parameters, even if physical displacements are similar to the localization error. The method is also robust with respect to deviations between data and model assumptions. Additionally, the method annotates the state probabilities at the single-molecule level, refines localizations (33) and extracts distributions of state durations. We demonstrate its versatility by analyzing two bacterial membrane proteins that diffuse slowly and transition rapidly between immobile and diffusive states.

## Results

### ExTrack is a maximum-likelihood method to detect different diffusion states in single-molecule tracks

We developed ExTrack, a maximum-likelihood estimation (MLE) method that contains two main modules: A fitting module fits a multi-state Markovian diffusion model to a data set of noisy single-molecule tracks. This module in-fers global model parameters including localization error, diffusion lengths, transition rates, and the initial fractions of molecules (at the first time point of all tracks). Part of these global parameters can also be provided by the user, and localization error can even be provided for each peak (35, 36) if desired. ExTrack is flexible in terms of the number of states and spatial dimensions. Additionally, it can explicitly consider molecules leaving the field of view, which otherwise introduces bias (17). Based on global parameters, a singlemolecule annotation module then estimates state probabilities *p*_*b*_(*i*) for molecules to reside in state *b* at each time point *i*. To characterize single-molecule tracks further, we devel-oped two additional modules: A position-refinement module refines molecule positions by taking advantage of spatial correlations between subsequent localizations, conceptually similar to (33). This feature allows to maintain high temporal resolution for state transitions, while attaining high spatial resolution for immobile molecules. A fourth module produces histograms of state durations. This module can reveal non-Markovian behavior, even if ExTrack assumes Markov transitions.

ExTrack is based on a Hidden Markov Model (HMM) that approximates a continuous-time process by a discrete-time Markov model (37). The method calculates the prob-ability density of observing each track given a set of global parameters (32). In principle, this calculation requires integrating the joint probability density of all single-molecule states, real positions, and observed positions over all possible sequences of hidden diffusive single-molecule states and over all possible real (physical) molecule positions (see Methods, section A). This problem is computationally intractable for long tracks (computational time scales as *N* ^*n*^, where *n* is the number of time points and *N* the number of states). To reduce computational time, we took advantage of the fact that the real position at a given time step is little influenced by the actual state a few time points away. This allows us to introduce a sliding-window approximation that reduces computational time to the order of *N* ^*m*+1^, where *m* is a user-defined window size (Methods, section A.4). We suggest *m* = 2− 7, depending on expected diffusion lengths.

In many HMMs it is assumed that state transitions can only occur at the time points of the measurement (37). However, this approximation introduces a bias towards higher fractions of fast diffusing molecules. Instead, we assume state transitions to occur at the middle of steps (see Methods, A.2). Additionally, ExTrack can consider sub-steps to further reduce bias at high transition rates.

ExTrack is available both as a Python library (38) and as a TrackMate module (39) on Fiji.

### Performance and comparison to alternative methods

First, we tested the performance of the ExTrack fitting module by applying it to computationally simulated noisy tracks of molecules (10.000 tracks of 10 positions each, if not stated otherwise) that transition between an immobile state (state 0) and a slowly diffusive state (state 1). The latter is characterized by a small diffusion length *d*_1_ = 2 *σ*, where *σ* is the localization error (Fig. 1a). The diffusion length is the typical physical displacement along each dimension: 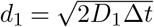 with *D*_1_ the diffusion coefficient and Δ*t* the time step. Here, we assume symmetric binding and unbinding rates *k*_u_ = *k*_b_ = 0.1 Δ*t*^−1^. Thus, on average, molecules reside in each state for ten time steps.

The dimensionless parameter 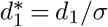 can be regarded as a signal-to-noise ratio. For a typical experiment, with *σ* = 20 nm and a time step of Δ*t* = 20 ms, a rescaled diffusion length of 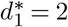 corresponds to a diffusion coefficient of *D*_1_ = 0.04 µm^2^.s^−1^, which is representative of typical membrane proteins *in vivo* (40, 41).

ExTrack reliably estimates all global model parameters (Fig. 1d) despite similar observed distances for immobile and diffusive molecules (Fig. 1c) and despite a low number of 10 localizations per track. Since molecules are considered at steady state in our example, the initial fraction of immobile molecules is given by *F*_0_ = *k*_b_*/*(*k*_u_ + *k*_b_).

Next, we simulated tracks for a three-state model, with 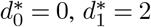 and 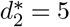, where transition rates are *K*_*i,j*_ = 0.1 Δ*t*−^1^ for all pairs of states. Fig. 1e demonstrates that ExTrack estimates global model parameters reliably. If the data contains long enough tracks, ExTrack can also correctly predict two immobile states of different lifetimes and their associated transition rates (Table S1). We will revisit more complex data sets below.

Returning to the simpler two-state model, ExTrack is capable to predict global model parameters reliably for a large range of model parameters (Fig. 1f). Predictions are accurate for diffusion lengths as low as the localization error and transition rates as high as 0.5 Δ*t*^−1^ (for independent variations of *k*_b_ and *k*_u_ see Fig. S1). To account for rapid transitions, we employed ExTrack considering two sub-steps. However, the method predicts parameters almost equally reliably without sub-steps (Fig. S2), while achieving improved computational time (Fig. S3).

Next, we compared ExTrack with the two MLE-based methods vbSPT (3) and anaDDA (42) that use absolute distances between localizations for parameter estimation. While vbSPT uses a HMM for the likelihood estimate (3) anaDDA is based on an analytical form of the distributions of apparent diffusion coefficients from short tracks (42). Both methods are restricted to a smaller parameter range than ExTrack (Fig. 1f-g and Fig. S4a) in the tested regime. The errors of parameter estimation by vbSPT are largely due to systematic bias, while the error of anaDDA is predominantly stochastic (Fig. S5) (42). We also tested a mean-field approximation based on track positions and considering hidden particle positions, the variational method UncertainSPT (32). We found that UncertainSPT performs worse and takes more computational time than ExTrack, anaDDA, or vbSPT (Fig. S4b).

### ExTrack is robust with respect to non-ideal motion properties

Single-molecule tracks in real cells often deviate from our basic model assumptions. Here, we investigated three different types of such deviations: i) variations of diffusion coefficients or localization precision, ii) finite track lengths due to a finite field of view or focal depth, and iii) physical confinement:

Diffusion coefficients can show intra- or inter-track variations (21, 43, 44), for example due to local variations of viscosity (43), and localization error can vary, for example if molecules are out of focus. We thus simulated tracks of a two-state model with diffusion coefficients or localization precision drawn from a chi-squared distribution with fixed mean and variable coefficient of variation (CV). First, we show that ExTrack gives very accurate predictions when localization error is specified for each peak instead of being treated as a single global fitting parameter (Fig. S6a). However, even when no prior information on localization error is given, ExTrack reliably predicts the average model parameters for variations up to 30-50% (Fig. 2a), in contrast to the distance-based methods anaDDA and vbSPT (Fig. S7). Track-to-track variations in diffusion coefficient of similar magnitude (up to about 50%) do also not affect predictions of average parameters (Fig. 2b).

**Figure 2.**
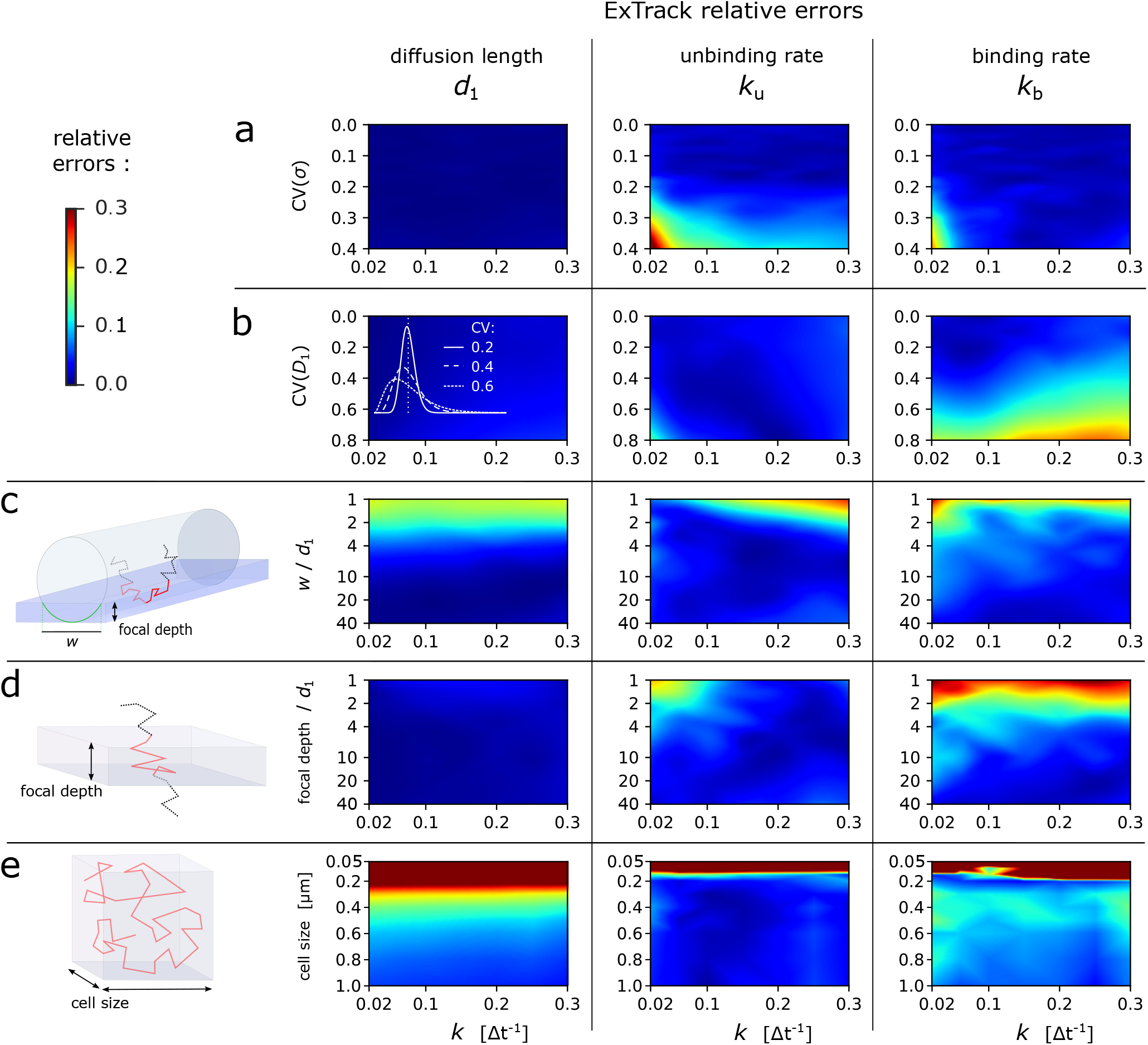
Robustness of ExTrack to various sources of bias. Heatmaps of relative errors on *d*_1_, *k*_u_ and *k*_b_ in case of two-state parameter fits to two-state simulations with one immobile state and one diffusive state, for different sources of bias as a function the source of bias (Y axis) and transition rates *k*. **a-b**: We simulated track-to-track variations of localization error *σ* (**a**) or diffusion coefficient *D*_1_ (**b**). Varied parameters followed chi-square distributions (white graphs in **b**) re-scaled so the mean localization error equals 0.02 µm (**a**) or the mean diffusion coefficient equals 0.25 μm^2^.s−^1^, which corresponds to a diffusion length of 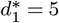 (**b**). **c**: Membrane proteins diffuse on a cylindrical surface and leave the field of view on the sides (see cartoon). We varied the width *w* of the field of view as indicated in the cartoon, while maintaining *d*_1_ and *σ* fixed. **d**: Cytoplasmic proteins can leave the focal plane anywhere (see cartoon). We varied the focal depth while maintaining *d*_1_ and *σ* fixed. **e**: Particles confined in a symmetric cube. We varied the box size while maintaining *d* and *σ* fixed. 10 replicates per condition. If not stated otherwise, *d*_0_ = 0 µm, *d*_1_ = 0.1 µm and *σ* = 0.02 µm. ExTrack settings: window length = 7, no sub-steps.

In situations, where the diffusion coefficient is even more broadly distributed, ExTrack can be used assuming a threestate model followed by aggregation of two diffusive states (Fig. S6b). We tested this aggregation approach with simulations of one immobile and five diffusive states, mimicking a broad distribution of diffusion lengths and jump distances (Fig. S6b, Table S2). The aggregated three-state approach reliably quantifies transitions between aggregated states and corresponding state fractions, thus providing a practical approach to the often encountered difficulty of choosing the right number of diffusive states.

Second, molecules can leave the field of view depending on microscopy modality and substrate geometry. For example, cytoplasmic molecules studied by confocal or epifluorescence microscopy diffuse in and out of the the focal plane, and proteins embedded or attached to a cylindrical membrane (for example, in bacteria) studied by TIRF microscopy leave the illumination field (Fig. 2c-d). Thus, immobile or slowly diffusing molecules are over-represented among long tracks, which has previously been described as ‘defocalization bias’ (17). We alleviate this bias by taking track termination into account explicitly (Methods, section A.6) similarly to previous approaches (17, 45). In both free 3D diffusion and diffusion along a cylindrical membrane, ExTrack reliably estimates model parameters as long as the typical dimension (focal depth or width of the field of view) is at least twice the diffusion length (Fig. 2c-d).

Finally, we tested the ability of ExTrack to analyse tracks of spatially confined molecules, as frequently found in membrane domains or small volumes such as bacteria or intracellular compartments. ExTrack performed robustly as long as confining dimensions are at least two to four times larger than the diffusion length (Fig. 2e).

### ExTrack computes state probabilities at every time point and refines positions

Next, we tested the performance of the single-molecule probabilistic annotation module of ExTrack, which is based on global model parameters and annotates state probabilities for every time point (31).

Fig. 3a-b shows tracks from the simulation of a two-state model with an immobile and a slowly diffusive state (*d*^∗^ = 2). Despite the small value of *d*^∗^, motion states are reliably estimated. High uncertainty is only observed at time points close to transition times (Fig. S8a), close to track boundaries, or if *d*^∗^ *<* 2 (Fig. 3c). To demonstrate the accuracy of the probabilistic annotation, we confirmed that among all molecules predicted to reside in the diffusive state with probability *p*_1_, the fraction of molecules actually diffusive also equals *p*_1_ (Fig. S8b). We also found ExTrack annotations to be robust with respect to wrongly chosen global parameters (Fig. S8c).

**Figure 3.**
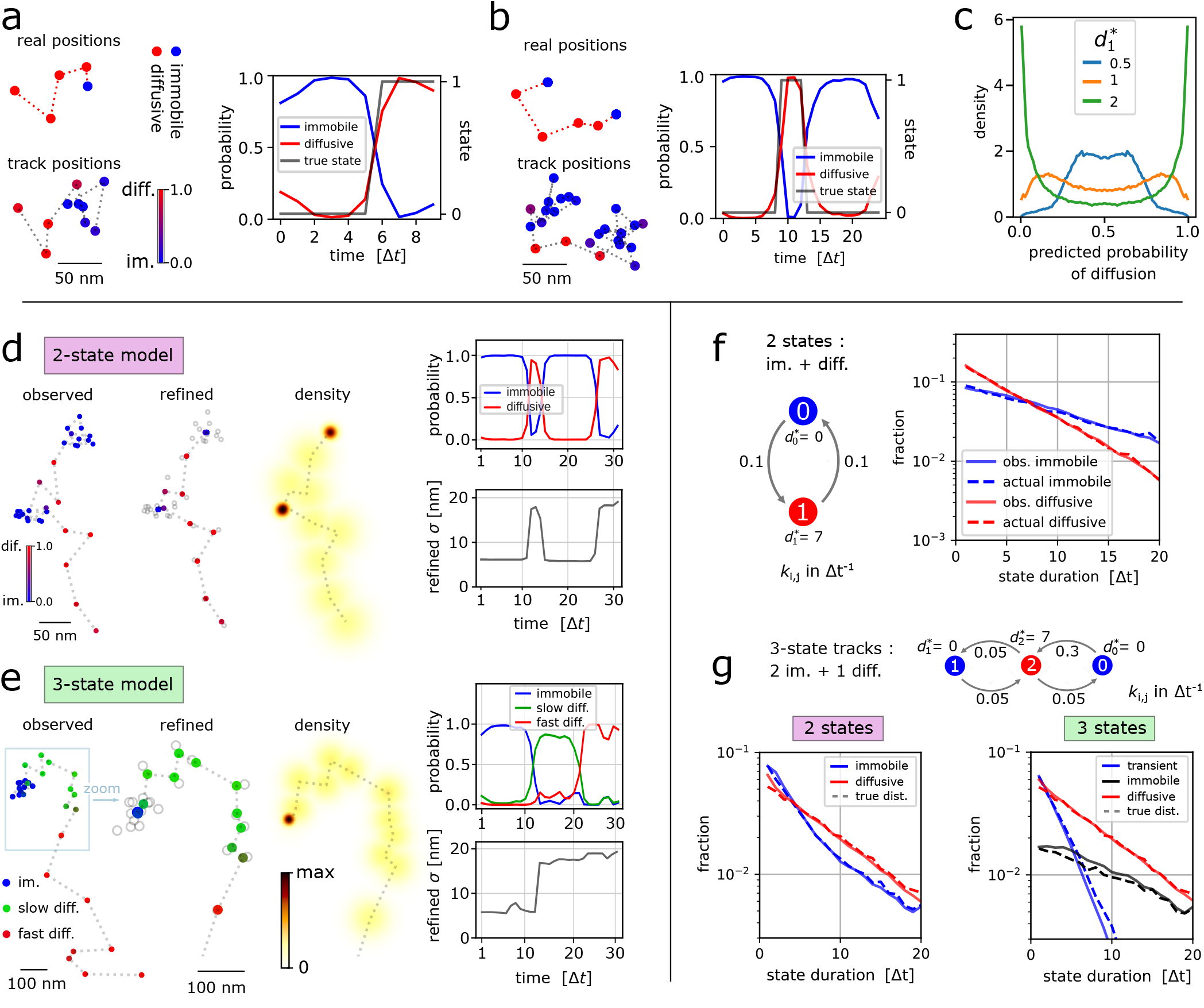
ExTrack annotates and refines single-molecule positions and extracts state-duration distributions. **a-b**: Example tracks from simulations of immobile and diffusive 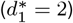 molecules with symmetric transition rates of 0.1 Δ*t*^*−*1^ with short tracks of 10 positions (**a**) or long tracks of 25 positions (**b**). Top: real positions with states in colors, bottom: noisy positions with probabilities (color bar), right: state probabilities along time. **c**: Distribution of the probability to be diffusive from similar simulations than in **a** for different 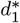. **d-e**: Position refinement module: Examples of 2-state track (**f**) and 3-state track (**e**). From left to right: Observed track and associated states probabilities; refined positions (colored) and observed positions in gray; probability density map of the consecutive positions. Top right: State probabilities as a function of time. Bottom right: standard deviation of the probability density of refined positions. Simulation parameters: 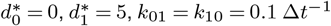 (**c**); 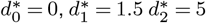 all *k* = 0.05 Δ*t*−^1^(**d**). **g**: Histogram module: state-duration histograms of tracks of at least 21 positions for the indicated 2-state model. Dashed lines: distributions from ground truth. **h**: Same as **e** for 3-state tracks with 2 immobile states. Left: ExTrack fit assuming a 2-state model; Right: ExTrack fit assuming a 3-state model.

While previous methods often classify molecules categorically into the most likely state (3, 32, 46) state probabilities allow discriminating regions of highly likely states from regions of intrinsically high uncertainty. However, even if categorically classifying molecules (Fig. S8d), ExTrack performs better than the binary method vbSPT (Fig. S8e). Next, we tested the capacity of ExTrack to refine positions by calculating the most likely physical position for each time point. Fig. 3d-e demonstrates that the position-refinement module effectively reduces the localization error of immobile molecules by 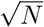, where *N* is the number of localizations in the immobile segment. This feature allows to obtain accurate positions of molecular binding sites inside cells, while still resolving state transitions dynamics.

### ExTrack computes distributions of state durations to characterize transition kinetics beyond the Markov assumption

ExTrack provides a histogram module that generates probability distributions of state durations. Instead of considering only the most likely set of states, ExTrack considers a large number of potential state vectors with their corresponding probabilities. To test the histogram module, we first simulated a Markovian two-state model. The predicted diffusive and immobile state durations are distributed exponentially, as expected, and in agreement with the simulated data (Fig. 3f). Therefore, any deviation from exponential decay can reveal more complex transition behavior: As an example, we simulated molecules that transition between two immobile states and one diffusive state (Fig. 3g). The histogram of immobile state durations then accurately reveals two sub-populations, even though ExTrack considers a two-state HMM model. Our approach thus indicates the presence of a third state, as confirmed by the exponential distributions of state durations after fitting the data to a three-state model (Fig. 3g).

The histogram approach is also relevant when the transitions are non-Markovian, for example if transition rates are spatially dependent (28, 47) or if states have minimum durations. In summary, the histogram module can help identify hidden or non-Markovian states and thus guide model choice.

### Application of ExTrack to experimental tracks of bacterial envelope proteins

To test our approach on experimental data, we used TIRF microscopy to track single green-fluorescent-protein (monomeric super-folder-GFP) fusions to two bacterial membrane proteins in *Escherichia coli*, each involved in one of the two major pathways of cell-wall synthesis.

First, we studied the cell-wall-inserting penicillin-binding protein PBP1b, which was previously described to reside in immobile or diffusive states (48, 49). However, transition rates and potentially hidden states remain unknown. When assuming a two-state model, ExTrack indeed reveals an immobile and a diffusive fraction (Fig. 4a), with the immobile fraction increasing with decreasing expression level (Fig. 4b) as expected (49). However, distributions of state durations obtained through the histogram module suggest the presence of at least two immobile populations with distinct unbinding rates (Fig. 4c). Since applying ExTrack assuming a three-state model revealed one immobile and two diffusive states (rather than two immobile states, Fig. S10), we also applied ExTrack with a four-state model (Fig. 4d-e, Fig. S11). The four-state model confirmed two diffusive states and two immobile states: among the immobile states we found a longlived state (lifetime of around 0.5 s) that is highly dependent on expression level (Fig. 4e), likely reflecting enzymatically active PBP1b, and a short-lived state with a lifetime of about 50 ms, likely reflecting non-specific associations with the cell wall. Thus, PBP1b displays rapid transitions between at least four different states.

**Figure 4.**
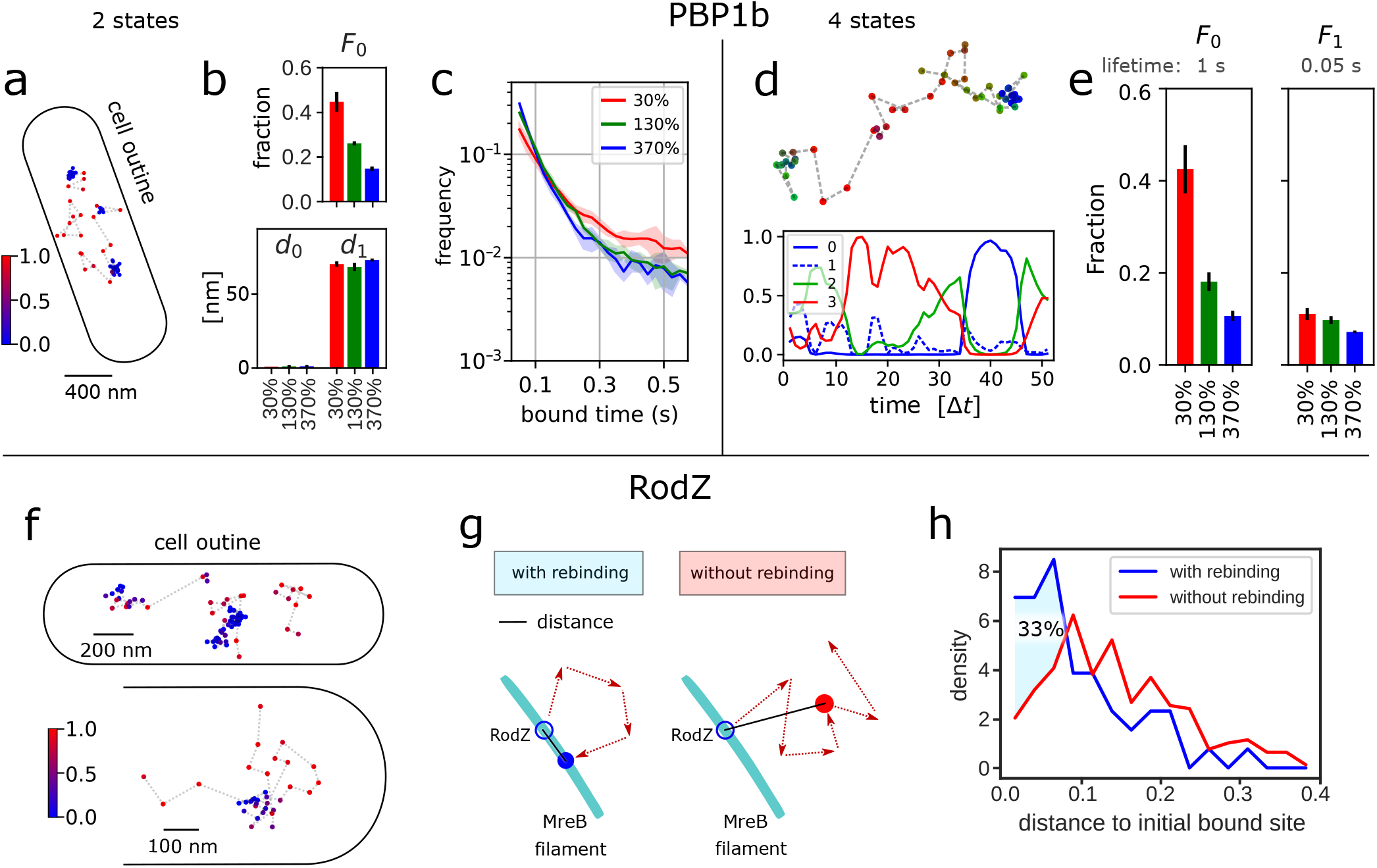
Characteristing PBP1b and RodZ motion. **a-e**: Analysis of GFP-PBP1b tracks (time step 25 ms) using ExTrack with two (**a-c**) or four states (**d**,**e**). **a**: GFP-PBP1b track (130% expression level). Color bar: Probability of diffusion. **b**: Diffusion lengths and fractions from two-state parameter fitting. (3 replicates, each with *>* 17.000 tracks of at least 3 positions, with average lifetimes from 6.3 to 7.5 positions). **c**: State-duration histograms of PBP1b tracks of at least 21 positions (*>* 600 tracks per replicate with average lifetimes from 28.5 to 32 positions), using global parameters from fitting a two-state model (**a** and Fig S9). **d**: Example PBP1b track (from 130% expression level) with associated state probabilities along time (first position on the left). **e**: Fractions from 4-state parameter fitting to the same datasets used in **a. f-h**: Analysis of GFP-RodZ molecules: **f**: RodZ tracks with overlapping binding sites. Color bar: Probability of diffusion. **g**: Cartoon illustrating rebinding of diffusive RodZ molecule to extended MreB-actin filament in two dimensions. Blue solid line indicates distance between initial position and position after four diffusive steps for molecules rebinding (left) or continuing diffusion (right). **h**: Histograms of distances between initial bound site and the site after 4 diffusive steps. Tracks rebind in closer vicinity to the initial binding site 33% more often than expected in case of random motion. Mann–Whitney U test: p-value = 1.6e-6. See Methods section H for details. Error bars and shaded regions: standard deviations between replicates.

Next, we investigated the motion of RodZ, a transmembrane protein that physically links cytoplasmic MreBactin filaments to a multi-enzyme complex that inserts new peptidoglycan while continuously moving around the cell circumference over minutes (50–52). Here, we studied the motion of GFP-RodZ on short time scales of seconds, where continuously moving complexes appear as immobile. Assuming a two-state model, the fitting module reveals that 70% of RodZ molecules are immobile (Fig. S13a-c), with a lifetime of about 0.7 s. This timescale is much shorter than the minute-long lifetime of the rod complex (13, 48, 50) demonstrating that a majority of immobile RodZ molecules is not stably associated. Instead, these molecules might transiently bind the MreB-actin cytoskeleton. Interestingly, RodZ molecules seem to often unbind and rebind in very close vicinity from the initial binding site (Fig. 4f, Fig. S14). Such behavior would be expected if RodZ could bind anywhere along extended MreB filaments, since filaments constrain diffusion in two dimensions (Fig. 4g). To test whether proximal rebinding occurs more often than randomly, we compared tracks that were initially bound, then diffusive for 4 steps, and then either rebound or remained diffusive (Fig. 4g). Short distances were indeed over-represented among rebinding molecules compared to molecules that remained diffusive (Fig. 4h). This behavior contrasts with PBP1b, which appears to bind to random sites (Fig. S13d). The annotation module of ExTrack thus allows us to identify spatial patterns of molecule binding that can be responsible for non-Markovian binding (28).

## Discussion

In summary, ExTrack provides a suite of robust tools to characterize single-particle tracks, extracting global model parameters, state probabilities at every time point, refined positions, and histograms of state durations, even if tracks are noisy, transitions are rapid, and tracks deviate from idealized model assumptions.

In tracking experiments, a major challenge is to identify the relevant number of immobile and diffusive states. Multiple previous methods obtain this number automatically (3, 19, 24, 33). However, at least some of these approaches tends to over-fit the data (25, 33). In more recent approaches, a high number of states is fixed followed by aggregation into one aggregated immobile state and one aggregated diffusive state, based on a user-defined diffusion-coefficient threshold (53). Here, we propose an alternative and iterative approach to complex tracking data: Data is initially fit to a coarsegrained twoor three-state model that can subsequently be expanded depending on desired variables and fitting results. For example, if one is predominantly interested in the exchange between immobile and diffusive molecules but not in the presence of multiple diffusive states, we propose a coarsegrained two-state or an aggregated three-state model that reliably predicts immobile-diffusive transitions, even if diffusion coefficient is variable or if molecules transition between different diffusive states (Fig. 2b, Fig. S6b). At the same time, ExTrack can also distinguish multiple diffusive states explicitly (Fig. 1e). Additionally, the distribution of immobile state durations can reveal the presence of multiple immobile fractions, which can then motivate the increase of the number of states.

The capacity of ExTrack to work with noisy single-molecule tracks is based on the explicit consideration of all sequences of states within a sliding window when computing the probabillity of every track, while states outside the sliding window are taken into account through averaging to limit computational time. In the future, this versatile principle can be extended to capture different and more complex dynamics, for example by considering persistent motion (6, 7), anomalous diffusion (54) and spatial maps of diffusion coefficients or states (44).

## Acknowledgements

We thank Andrey Aristov for setting up the TIRF microscope, Antoine Vigouroux for the GFP-PBP1b strains, and Gizem Özbaykal for her guidance for experiments. We also thank Felipe Bendezú and Piet De Boer for the GFP-RodZ fusion. This work was supported by the European Research Council (ERC) under the Europe Union’s Horizon 2020 research and innovation program [Grant Agreement No. (679980)] to SVT, the French Government’s Investissementd’Avenir program Laboratoire d’Excellence “Integrative Biology of Emerging Infectious Diseases” (ANR-10-LABX62-IBEID, SVT), France BioImaging (ANR-10-INBS-04, JYT), the Mairie de Paris “Emergence(s)” program to SVT, a NSERC Discovery Grant to SVT, a FRQS Salary Fellowship to SVT, as well as support from the Volkswagen Foundation to SVT.

## Methods

In the following sections we will describe the ExTrack method with its four different modules: the fitting module (Section A), the annotation module (Section B), the position-refinement module (Section C) and the histogram module (Section D). Subsequently, we will describe the generation of computationally simulated datasets (F), the interpretation of results by vbSPT (Section G), and the experimental methods (Sections H to J).

### A. ExTrack Fitting Module

#### A.1. Introduction

ExTrack fits a multi-state diffusion model to noisy single-particle tracking data. We assume that tracks come about according to a continuous-time Markov model, where molecules transition randomly between *N* diffusive states at rates *k*_*i,j*_. As long as molecules reside in state *i*, they undergo Brownian diffusion with diffusion coefficient *D*_*i*_. Additionally, observed positions *c*_*i*_ are displaced from real positions *r*_*i*_ according to a Gaussian distribution *f*_*σ*_(*c*_*i*_− *r*_*i*_), where the standard deviation equals the localization error *σ*. Here and in the following, *f*_*x*_(*y*) generally denotes a Gaussian distribution of standard deviation *x*. The *N* -state diffusion model is thus characterized by the parameters *θ* = (*σ, D*_*i*_, *F*_*i*_, *k*_*i,j*_) for all states *i, j* ∈1, …, *N*. Here, *F*_*i*_ are the fractions of molecules residing in state *i* at the first position of the track. Later, we will introduce additional parameters for additional spatial dimensions and for the treatment of non-constant track lengths.

Parameters are estimated based on a maximum likelihood estimate approach (MLE), which, in turn, is based on accurately computing *f*_C_(*C θ*), the probability density of observing a track of positions *C* = (*c*_1_, *c*_2_, …, *c*_*n*_). The likelihood of the parameters given the data *L*(*θ*| all *C*) then equals the product of *f*_C_(*C* |*θ*) for all tracks *C*. By maximizing this function, we can find *θ*^∗^, the optimal estimator of the underlying parameters. Optimal parameters *θ*^∗^ are found by MLE using the Powell method. ExTrack also allows to fix individual or multiple parameters. This generally speeds up the fitting process and reduces variations in the remaining parameters. In this realm, we also found that fixing the localization error to a slightly wrong value has little impact on the fitting of the other parameters as long as it does not deviate by more than about 20 −30%. Here and in the following we treat localization error as a model parameter, but the user can also provide spot-specific localization errors based on photon counts (35).

In the following sections, we will first compute *f*_C_. This calculation is presented in one spatial dimension (1D). However, the model is easily extendable to 2D or 3D due to the independence of the displacements and localization error in each axis, as we will see below.

#### A.2. Parameter fitting based on the probability distribution of observed positions

Tracks are generally described by their sequence of observed positions *C* = (*c*_1_, *c*_2_, …, *c*_*n*_). Those positions come about based on the sequence of physical molecule positions *R* = (*r*_1_, *r*_2_, …, *r*_*n*_), which, in turn, are the stochastic result of the sequence of diffusive states *B* = (*b*_1_, *b*_2_, …, *b*_*n*_).

For a given track *C*, the probability density function *f*_*C*_ can be calculated from *f*_C,B,R_, which is the joint probability density function of observed positions *C* = (*c*_1_, …, *c*_*n*_), real (physical) molecule positions *R* = (*r*_1_, …, *r*_*n*_), and timedependent diffusion states *B* = (*b*_1_, …, *b*_*n*_), by integration over all possible values of *R* and by summation over all possible values of *B*:

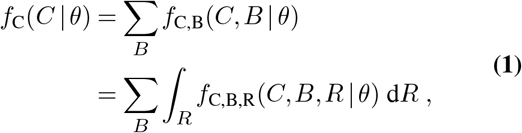

where we defined the joint probability density function *f*_C,B_(*C, B* |*θ*) of having *C* and *B* given *θ*. The joint probability density *f*_C,B,R_ can be decomposed into a product of three terms: the a priori probability of *B*, the probability density of the physical displacements *f*_R|B_, and the probability density of the distances between real position and observed positions *f*_C|R_, respectively:

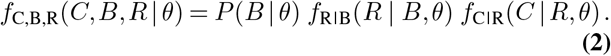

Here, the a priori probability of the sequence of states *P* (*B* |*θ*), which we refer to as *β* for brevity, results from the Markovian processes of transitioning between states (55). *β* is obtained as

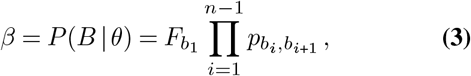

where 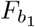 indicates the fraction of molecules in state *b*_1_ at time point 1, and where 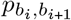 indicates the probability to transition from state *b*_*i*_ at time point *i* to *b*_*i*+1_ at time point *i* + 1. The transition probabilities can be computed from the continuous-time transition rates (see Subsection A.5). The initial fraction 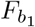 can either be an independent parameter or constrained by transition rates at steady state (for a twostate model, *F*_0_ = *k*_10_*/*(*k*_10_ + *k*_01_)). *f*_R|B_ is the probability density function of real positions *R* given the sequence of states *B* and *θ. f*_C|R_ is the probability density function of the sequence of observed positions *C* given the real positions *R* and *θ. f*_C|R_ can be expressed as a product of Gaussian distributions with standard deviation equal to *σ*:

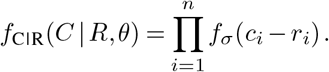

Next, we express *f*_R|B_(*R* | *B, θ*) as a product of Gaussians:

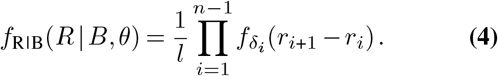

Here, *l* is the length of the space of real positions. Without any prior on *R*, we consider the limit *l*→∞. However, since *l* only appears as a constant prefactor, we can ignore it in the calculation of the log likelihood. According to previous suggestions, the width of the distribution *δ*_*i*_ should equal the diffusion length corresponding to the current state *b*_*i*_ (37). However, this discretization of the continuous-time Markov process introduces a bias towards diffusive motion. This is easily illustrated in case of an immobile-diffusive model: There, a particle initially immobile starts moving before the first measured time point where the particle is observed to be diffusive. Similarly, it stops moving after the last time point, where it is observed to be diffusive. Then, a model assuming diffusion to only be dependent on the current hidden state will overestimate the time spent in diffusive state by up to half a step in case of high diffusion. Logically, this results in underestimating the binding rate (Fig. S4c), immobile fraction and diffusion length when transitions are frequent. To alleviate this issue, we assume transitions to occur at the middle of two time points. The standard deviation of the probability density function 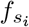 then equally depends on states at each of the two subsequent time points, with

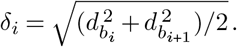

This assumption effectively decreases the bias inherent to the discrete approximation of continuous tracks (Fig. S4c). Later, we will also introduce sub-steps between time points that improve the approximation (see Subsection A.5).

Taking advantage of the expressions of *f*_C|R,*θ*_ and *f*_R|B,*θ*_, Equation (2) becomes

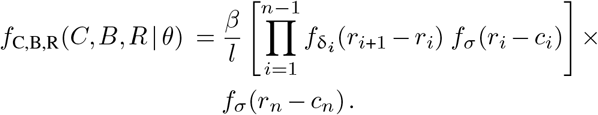

Inserting this expression into Equation (1), we then integrate step-wise over all real positions *R* = (*r*_1_, …, *r*_*n*_). This allows us to use the recusion principle (30) to compute *f*_C,B_(*C, B* | *θ*): The first step consists in integrating the two Gaussian distributions dependent on *r*_1_ (displacement and localization error terms). This integration results in a Gaussian distribution 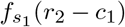, of standard deviation 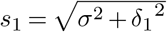 (constituting a convolution of two independent random variables with Gaussian distributions). For each of the next integrals over *r*_*i*_, we integrate the product of three Gaussian distributions (for the random displacement *r*_*i*+1_ − *r*_*i*_, the localization error *r*_*i*_ − *c*_*i*_, and the previous term of the distribution 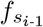). The result of this integration can be expressed by a scalar *K*_*i*_ times a Gaussian distribution 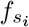, according to

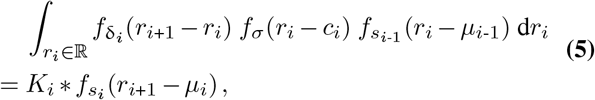

where 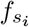 is a Gaussian distribution of standard deviation *s*_*i*_ and mean 0. The standard deviation *s*_*i*_ and mean *µ*_*i*_ can be expressed depending on *s*_*i*-1_ and *µ*_*i*-1_:

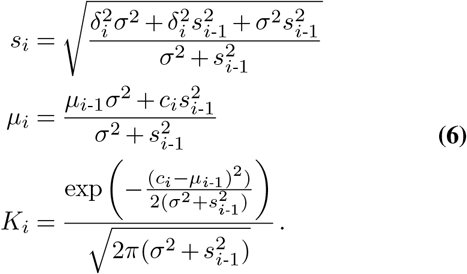

The recursion process can then be summarized by the sequences *s*_1_ : *s*_*n*-1_ and *µ*_1_ : *s*_*n*-1_ which depends on *C,B* and *θ*. At the last step (integration over *r*_*n*_), we integrate the product of the two remaining Gaussian distributions: the previous term 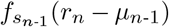 and the localization error term *f*_*σ*_(*c*_*n*_ − *r*_*n*_), as described for the first step to compute the density function *f*_C,B_(*C*, | *B θ*).

Finally, we compute the value of the probability density function of the observed track *f*_C_(*C* | *θ*) as the sum of *f*_C,B_(*C, B* | *θ*) over all possible *B*.

#### A.3. Extension to 2D and 3D

Since diffusive motion is independent in each spatial dimension, the principle described above for one dimension can simply be extended to two or three dimensions by multiplication of independent distribution functions. For example, in 2D, the function *f*_C,B_(*C, B* | *θ*) is simply replaced by the product *f*_C,B_(*C*_*x*_, *B* | *θ*)*f*_C,B_(*C*_*y*_, *B* | *θ*).

In principle each axis can have a different localization error and different diffusion lengths for each state. This is especially true for localization in the direction of the optical axis compared to the lateral axes. ExTrack therefore allows to have independent localization errors for each axis.

Alternatively, the user can also provide localization error for each peak, for example using the Cramer-Rao lower bound estimate (32, 56). Since these and other estimators might underestimate the true localization error, peak-wise localization estimates can also be implemented as scaling factors that are then assumed to be linearly related to the true localization error estimated by ExTrack.

#### A.4. Using a window to reduce calculation time

This method has a number of operations which initially scales with *N*^*n*^, where *N* the number of states and *n* the number of time points. This means the calculation time can become unrealistically long when analysing long tracks. To alleviate this issue we developed a window method to allow it to work with longer tracks in a reasonable time scaling with *nN*^*m*+1^ (*m* the window length of minimal value 1). For computational reasons, we advise to use a window length of 7 for 2-state models, 5 for 3-state models and 3 or 4 for more states.

Here, we briefly motivate and describe the implementation of the window method: During the recurrence process described above, Equations (5) and (6), 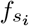 can be regarded as a density probability function of the position *r*_*i*+1_ knowing the previous observed positions *c*_1_, …, *c*_*i*_ and states from positions 1 to *i* + 1. We realized that the current localization of a particle is very little affected by its state *m* steps ago when *m* »1. Thus, for a given track, two sequences of states varying only for their first state should give very similar 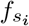. The values *µ*_*m*_ and *s*_*m*_ of these normal distributions should also be similar.

For example, if the track has been diffusive during at least one of the positions from steps *i − m* to *i*, the current observed position *c*_*i*_ is much more informative for the real position *r*_*i*+1_ than the first observed position *c*_1_. If the molecule has been immobile from position *i* − *m* to *i*, all observed positions *c*_*i*-*m*_ to *c*_*i*_ are equally informative. However, even the past 5-7 positions are likely sufficient to predict the distribu-tion 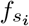.

As we saw in Subsection A.2, for a given sequence of states *B*, computing *f*_C,B_(*C, B* | *θ*) is nothing but computing three sequences *s*_1_:*s*_*n*_, *µ*_1_:*µ*_*n*_ and *K*_1_:*K*_*n*_ until the last step where we simply have to compute 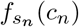.

In the recursion process, in case of a two-state model, we start by computing four values each for *s*_2_, *µ*_2_ and *K*_2_ that we will differentiate as 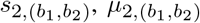 and 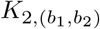, corresponding to the transition between the state *b*_1_ at time point 1 and state *b*_2_ at time point 2. For this recursion step, the four values of *s*_2_, *µ*_2_ and *K*_2_ arise from the following four combinations of states (0, 0), (0, 1), (1, 0), (1, 1). At *i* = 3, we get 8 possible state combinations, at *i* = 4 we get 16, etc. At step *m*, any sequence of states has a characteristic 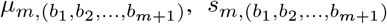 and 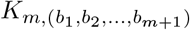. In order to limit the number of consid-ered sequences to 2*m*, we can merge *µm*,(0,*b*_2_,…,*b*_*m*+1)_ and *s*^2^):*µm*,(1,*b*_2_,…,*b*_*m*+1_) to an average *µm*,(∗,*b*_2_,…,*b*_*m*+1_) (same for 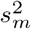):

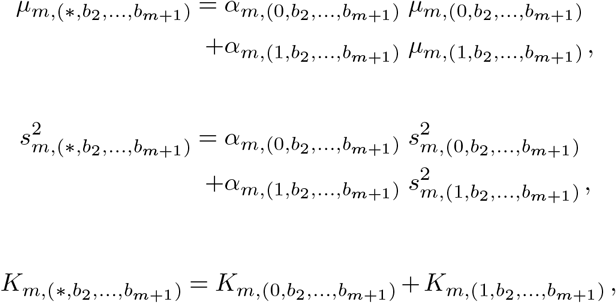

where *α* are the averaging weights according to the joint probability density

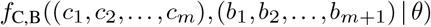

of observed positions *c*_1_, …, *c*_*m*_ and states *b*_1_, …, *b*_*m*+1_. For brevity we express this probabillity density as *Pm*,(*b*_1_,*b*_2_,…,*b*_*m*+1_) in the following expression for *α*:

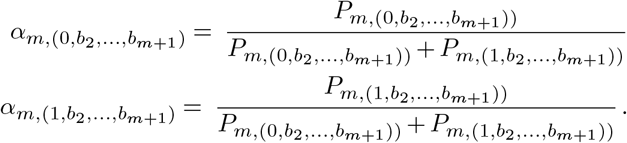

In this way, two sequences of states are merged (for example, the sequences starting with (0,0,0,1,1,1) and (1,0,0,1,1,1)). We thus reduce the number of *µ*_*m*_, 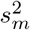 and *K*_*m*_ from 2*m*+1 to 2*m*. By recursion of this principle over all steps from *m* to *n* − 1 we limit the computation time to 2*m*+1 for a two-state model, or, more generally, to *N*^*m*+1^ for a *N* state model. In the following Subsection A.5, we will introduce sub-steps between the discrete observation time points. Our approach is easily generalized to sub-steps by considering state vectors.

Applying our approach with a window length *m* = 5 − 7, we observed similar functional dependencies of the likelihood on the model parameters *θ*, allowing us to drastically speed up our method without loosing accuracy.

#### A.5. Approximating continuous transitions with a discrete model of one or multiple sub-steps per time frame

ExTrack fits data of a continuous-time process to a discrete-time Markov model. Without the introduction of sub-steps, ExTrack assumes that transitions can only happen once per time step. It then estimates transition probabilities per time step, which must be translated into transition rates *k*_*i,j*_ that describe the continuous-time Markov model. For continuous-time Markov processes, transition probabilities can be converted into rates according to a simple relationship *P* = *e*^*G*Δ*t*^, where *P* is the transition probability matrix, which contains the transition probabilities *p*_*i,j*_ from state *i* to *j*, and where *G* is the generator matrix with elements *G*_*i,j*_ = *k*_*i,j*_ for 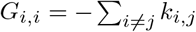 (57). Here, the transition probabilities *p*_*i,j*_ allow the molecule to transition from state *i* to *j* via any number of intermediate states.

However, the implementation of this relation into ExTrack leads to a systematic overestimation of transition rates. The reason for this overestimation is found in our approximate representation of the distribution of physical displacements between time points (Equation (4)), which is based on the false assumption that transitions can only occur at the middle of steps, contrary to the continuous-time nature of the underlying physical process. We found that this error could be compensated for by using a slightly different approximation for state transitions *p*_*i,j*_ = 1 − exp(− *k*_*i,j*_ Δ*t*). In the limit of small *k*_*i,j*_Δ*t*, this approximation asymptotically equals the exact expression *P* = *e*^*G*Δ*t*^, which also asymptotically equals *p*_*i,j*_ = *k*_*i,j*_Δ*t*. We found that ExTrack using the approximate relationship performs better in the case of two-state and three-state models for a large range of transition rates. However, ExTrack (python version) also allows using the generator-matrix based relationship, if the user desires.

When transition rates are high (when *k* Δ*t >* 0.4 for the two-state model), our method allows to subdivide time steps into a number of *u* sub-steps (where *u* = 2 corresponds to dividing each step into two). This allows ExTrack to account for multiple transitions and transition times that are different from the midpoints of time steps Δ*t*. To take states at sub-steps into account, we introduce a new state vector *B* = (*b*_1,1_, *b*_1,2_, … *b*_1,*u*_, …, *b*_*n*-1,*u*_, *b*_*n*,1_)) and new physical positions, that require integration according to Equation (1). This integration is straight-forward: The probability *β*, (Equation (2)) is simply replaced by the product of all state transitions between subsequent sub-positions. The probability distribution of real positions (Equation (4)), is replaced by the corresponding distribution of sub-positions. Since the physical displacement during Δ*t* is the sum of Gaussian random variables (the sub-displacements), the functional form of Equation (4) (a product of *n* − 1 Gaussians) can be maintained while replacing the standard deviations 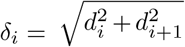 by 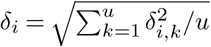, where *δ*_*i,k*_ are the corresponding diffusion lengths for the sub-steps.

The use of a window length *m* will allow the user to do accurate computations for *m* sub-steps. Thus, at a given *m*, the number of observed positions *c* considered within the window equals floor(*m/u*). To consider the same amount of observed positions per window, one thus needs to increase the window length. A trade-off between number of states, window length, and number of sub-steps has to be found (see Subsection E).

#### A.6. Extension of ExTrack to consider a finite field of view

Tracks can terminate due to different reasons: photobleaching, diffusive molecules leaving the field of view, or molecules transiently not being detected. The process of leaving the field of view requires diffusive motion. Observation of long-lived molecules within a finite field of view can thus show a bias towards non-moving or slowly moving molecules. An extension of ExTrack can take this bias into account by explicitly modeling the probability of track termination. We consider two contributions to track termination: first, a constant termination probability *p*_*K*_, which is independently of the motion state. This probability summarizes photobleaching and the probability to not detect a molecule, for example because of low signal to noise ratio; second, a probability of leaving the field of view (or observation volume) *p*_*L*_ that depends on the diffusion length and the dimensions of the field of view. In case of a cytoplasmic particle tracked through epi-fluorescence or confocal microscopy, the monitored length is the depth of field (or focal depth). In case of a membrane protein moving around a cylindrical cell imaged in TIRF microscopy, the monitored length is a fraction of the cell diameter (Fig. 2c-d).

In principle, *p*_*L*_ can be calculated depending on the position of the molecule with respect to the boundaries of the field of view. However, we decided to implement an approximate form of *p*_*L*_(*δ*_*i*_) that does not require this information and instead considers the position of the observed molecule as random inside the field of view. Within this approximation, the probability of leaving the field of view is given by

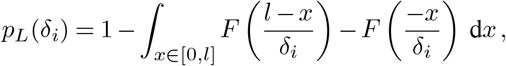

where *F* (*x*) is the cumulative density function of the standard normal law.

We thus modify *f*_C,B,R_(*C, B, R* | *θ*) in Equation (2) by multiplication of the left-hand terms with the probability of observing a track of *n* positions, which is given by

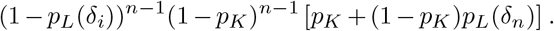

### B. Annotation module

The annotation module allows to compute the probabilities to be in any state at any time point of all tracks. According to conditional probabilities and results from Section A we can compute the probability of the sequences of states having the parameters *θ* for each track *C*:

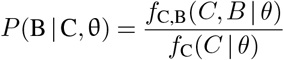

For a given track *C*, at each time point *i*, the probability of the current state *b*_*i*_ to be in state *s* ∈ { 0, 1} can then be computed by summing over all *B* with *b*_*i*_ = *s*:

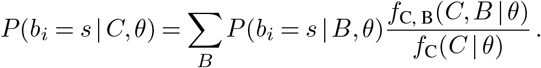

The annotation module can also take advantage of the window approximation decribed in Subsection A.4 to reduce computational time and make the computation tractable in case of long tracks.

Since the annotation module does not require parameter fitting and thus many iterations, the window length can be chosen larger than for the fitting module. A large window length is also more important for precise state prediction than for accurate global parameter fitting.

### C. Position-refinement module

ExTrack can improve the estimation of molecule positions based on a track, in particular if molecules move slowly. Positions can be estimated by computing the probability density function of each real positions *r*_*i*_ (32). To do so, we compute *f*_C_(*C* | *θ*) without integrating at position *r*_*i*_. This results in a probability density function *f* (*r*_*i*_ *C*, | *θ*) (Fig. 3d-e) which is a sum of Gaussian functions for each sequence of states. While this probability density can be obtained explicitly, it is much faster to obtain the expected value and standard deviation of the density function. Those values are computed by averaging the parameters of the Gaussian distributions associated with each sequence of states weighted by their respective probability. Like for the fitting method, the window method is applied (Subsection A.4).

### D. Histogram module

Computing state-duration histograms allows to assess non-Markovian transition behaviors or to reveal multiple hidden states with different transition rates. For a given state, the resulting rate is then the sum of track-termination rate (bleaching, track termination due to low SNR, leaving the focal plan) and the transition rates to other states. If the track-termination rate is low, the histogram allows to identify one or multiple transition rates (see, for example, Fig. 3f-g). Picking only long tracks can help removing the contribution from bleaching. ExTrack estimates the histograms *h*_*s*_ for each state *s*:

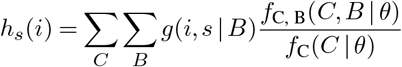

with *g*(*i, s* | *B*) the number of sequences of *i* consecutive time points of state *s* in the sequence of states *B*. As long tracks have to be assessed and all states of *B* kept in memory, the window method cannot be applied, we then only keep the most likely *B* (1000 in Fig. 3f-e).

### E. Implementation and computational time

ExTrack is available as a Python (Python 3) package (38). A version with the core functionality is also available as a TrackMate module (39) on Fiji. The TrackMate implementation can fit data to a two-state model and annotate states according to the results from the fit or manually chosen parameters. It then allows interactive visualization of tracks colored with state probabilities for each displacement. We allow parallelization with GPU (cupy library) (python version only) or multiple CPUs (both python and TrackMate versions).

As mentioned in Subection A.5, a trade-off between number of states *N*, window length *m*, and the number of substeps *u* has to be found for reasonable computational time. When running the ExTrack fitting module on a computer with Intel® Core™ i7-9700 processor (10 000 tracks of 10 positions) for 200 iterations using a two-state model, a window length of *m* = 2, and no sub-step (*u* = 1) the analysis can be as fast as 20 seconds. For the dependency of computational time on numbers of states, sub-steps, and window length see Fig. S3.

To save computational time, we recommend to initially run ExTrack with low values of *u* and *m* and then to increase *u* if model predictions suggest high transition rates or *m* for low predicted diffusion lengths. Specifically, we suggest to make the following adjustments: If localization error is negligible, for instance if there is no immobile state and all *d*_*i*_ *>* 2*σ*, window length *m* can be set to its minimal value of 1. Similarly, *m* = 1 should perform alright when there is one immobile state and all diffusive states have large *d*_*i*_ *>* 5). In such cases, multiple sub-steps can be used at little computational cost. More generally, if predicted transition rates are larger than 0.4 Δ*t*^-1^ but localization error is not negligible, we suggest increasing *u* to 2 for most accurate estimates (Fig. S2 vs Fig. 1f). In the hardest cases of small *d*\* ≲ 2 and high transition rates (;2 0.4), we recommend using *u* = 2 and *m* ≥ 8.

### F. Computational simulation of tracks

To test the predictive power of the different methods, we conducted overdamped Brownian Dynamics simulations of tracks in two or three spatial dimensions with molecules transitioning randomly between the different states at discrete time points. To mimic a continuous-time Markov model for state transitions we used a small time step *τ* = Δ*t/*50 « 1*/k*_*i,j*_, where *k*_*i,j*_ are the transition rates. Brownian Dynamics simulations were carried out by randomly drawing physical displacements in each spatial dimension from Gaussian distributions of standard deviation 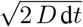, where *D* is the diffusion coefficient corresponding to the diffusive state. An additional Gaussian distributed noise of standard deviation *σ* was added to simulate localization uncertainty.

Except if specified otherwise, we simulated 10 000 tracks of 10 time points with localization error *σ* = 0.02 µm, Δ*t* = 0.06 s, *d*_0_ = 0 µm, *d*_1_ = 0.1 µm, bound fraction *F*_0_ = 0.5, transition fractions per steps *k*_01_ = *k*_10_ = 0.1 Δ*t*^-1^, with infinite field of view and perfectly stroboscopic tracks. We also assumed that molecules reached steady state, that is,

To test the robustness of ExTrack and the other methods to more complex behaviors we also simulated tracks with variations of localization error or diffusion coefficients, a finite field of view or physical confinement as follows (see Fig. 2 for illustrations):

Track-to-track variations of localization error (or diffusion coefficients) were simulated with localization error *σ* (or diffusion coefficient *D*) following *χ*^2^ distributions of given coefficients of variation and mean 0.02µm (or 0.25 µm2.s-1 for *D*), Δ*t* = 0.02 s. Models with multiple diffusion states were simulated as continuous-time transitions with model parameters detailed in Table S2.

To simulate a finite field of view in two dimensions, we simulated tracks in a box that is infinite in one spatial dimension (*y*) and finite in the other dimension (*x*) with size 3*l*, where *l* is the size of the field of view. All tracks or part of tracks that fall into the field of view are considered for further analysis. A single particle can thus result in several tracks if leaving the field of view and coming back. A finite field of view in three spatial dimensions was simulated analogously: The simulation box is infinite in *x*- and *y*-directions, while the box has periodic boundary conditions in the *z*-direction.

To simulate physical confinement, we considered tracks to move within a square area of indicated side length, using reflecting boundary conditions.

### G. Comparison to vbSPT

To compare our results with vbSPT we fixed the number of states to two so both algorithms performed exactly the same task. vbSPT does not consider localization error but a metric that we will call *u*. In case of pure diffusion, *u* = *D* Δ*t* but in case of immobile particle with localization error *u* = *σ*^2^*/*2 in principle. We can thus infer *σ* and *D* according to 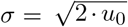 and *D* = (*u*_1_ − *u*_0_)*/*Δ*t*.

### H. Computational analysis of molecule rebinding

To assess the propensity of RodZ molecules to rebind in close vicinity of their initial binding site, we first annotated tracks using parameters obtained from the ExTrack fitting module. We considered the 16 first time points of tracks of at least 16 time points. Among tracks labeled as initially immobile for at least 3 time points (*p*_immobile_ *>* 0.5) then diffusive for 4 time points (with at least three time point of probability *p*_diffusive_ *>* 0.7), we grouped tracks into two subgroups, the ones rebinding right after and the ones, which continue to diffuse for at least 1 more time point. The histograms represent the distributions of distances between initially bound position and the position at the 4th time point after unbinding. If molecules were to rebind at random locations, the distributions of the distribution of distances for rebinding particles should be the as for particles, which continue to diffuse. Neither PBP1b (Fig. S13d) nor tracks obtained from immobile fluorescent beads (of similar signal-to-noise ratio) showed any significant rebinding, which excludes wrong conclusions on RodZ data due to miss-annotations.

### I. Cell cultures

We used the IPTG-inducible GFP-RodZ strain FB60(iFB273) (Δ*rodZ*, Plac:: *gfp-rodZ*) by (58) and the GFP-PBP1b-containing strain AV51 (*msfgfp-mrcB*, Δ*mrcA*) (49). Cells were grown overnight at 37°C (shaking) in LB medium and then washed and diluted at least 1:1000 in M63 minimal medium (Miller, 1972) supplemented with 0.1% casamino acids, thiamine (5 × 10-5 %), glucose (0.2%) and MgSO4 (1 mM) and grown for 6 hours to early exponential phase (maximum OD600 of 0.1) at 30°C (shaking). Cells were then spread on an agar pad made from the same M63 media as described above. RodZ production was induced with 100 *µ*M IPTG. In the strain AV51, CRISPR repression of msfGFP-PBP1b is induced with 100 ng/ml of anhydro-tetracycline (Acros Organics). When necessary, strains were supplemented with kanamycine (50 *µ*g/ml) or carbenicillin (100 *µ*g/ml) during overnight cultures. Biological replicates result from independent cultures starting from separate colonies.

### J. Single-particle tracking of msfGFP-PBP1b and GFP-RodZ proteins

Cells were all positioned in the same focal plan in between an agar pad (1%) and a coverslip to be imaged in TIRF microscopy. Coverslips were cleaned by 60 min sonication in saturated KOH solution followed by two washing steps (15 min sonication in milli-Q water). Single-particle tracking of GFP-PBP1b was performed with a custom-designed fluorescence microscope based on an ASI Rapid Automated Modular Microscope System, equipped with a 100x TIRF objective (Apo TIRF, 100x, NA 1.49, Nikon), Coherent Sapphire 488-200 laser, and a dichroic beamsplitter (Di03-R488/561-t3-25 × 36, Semrock). Excitation was controlled with an acoustooptic tunable filter (AA Optoelectronics) through an Arduino (15 ms light exposure per frame). Images were acquired using an Andor iXon Ultra EMCCD camera with an effective pixel size of 130 nm. Image acquisition was supervised with MicroManager.

Data analysis with ExTrack was restricted to tracks with at least 3 position. For long tracks, only the first 50 positions were analyzed.

## Supplemental Figures

**Supplementary Figure 1.**
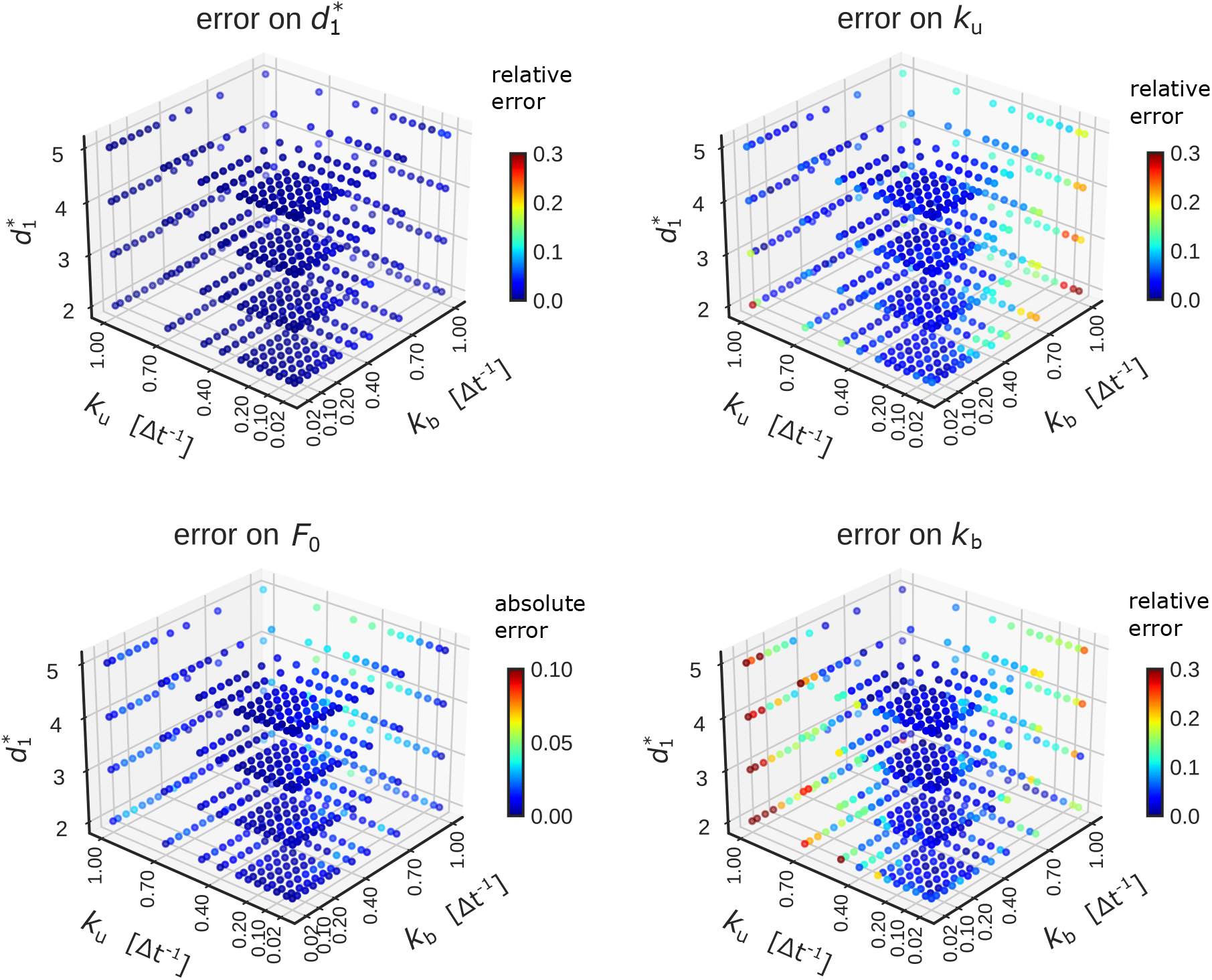
ExTrack parameter fits for independently varied binding and unbding rates. 3D map of the mean error on extracted parameters from simulations similar to those in Fig. 1, of two-state models with one immobile and one diffusive state as a function of diffusion length 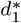, unbinding rate *k*_u_ and binding rate *k*_b_. Errors are obtained from 5 replicates. Errors are indicated as absolute or relative errors, as indicated. ExTrack settings: no sub-steps, window length = 7.

**Supplementary Figure 2.**
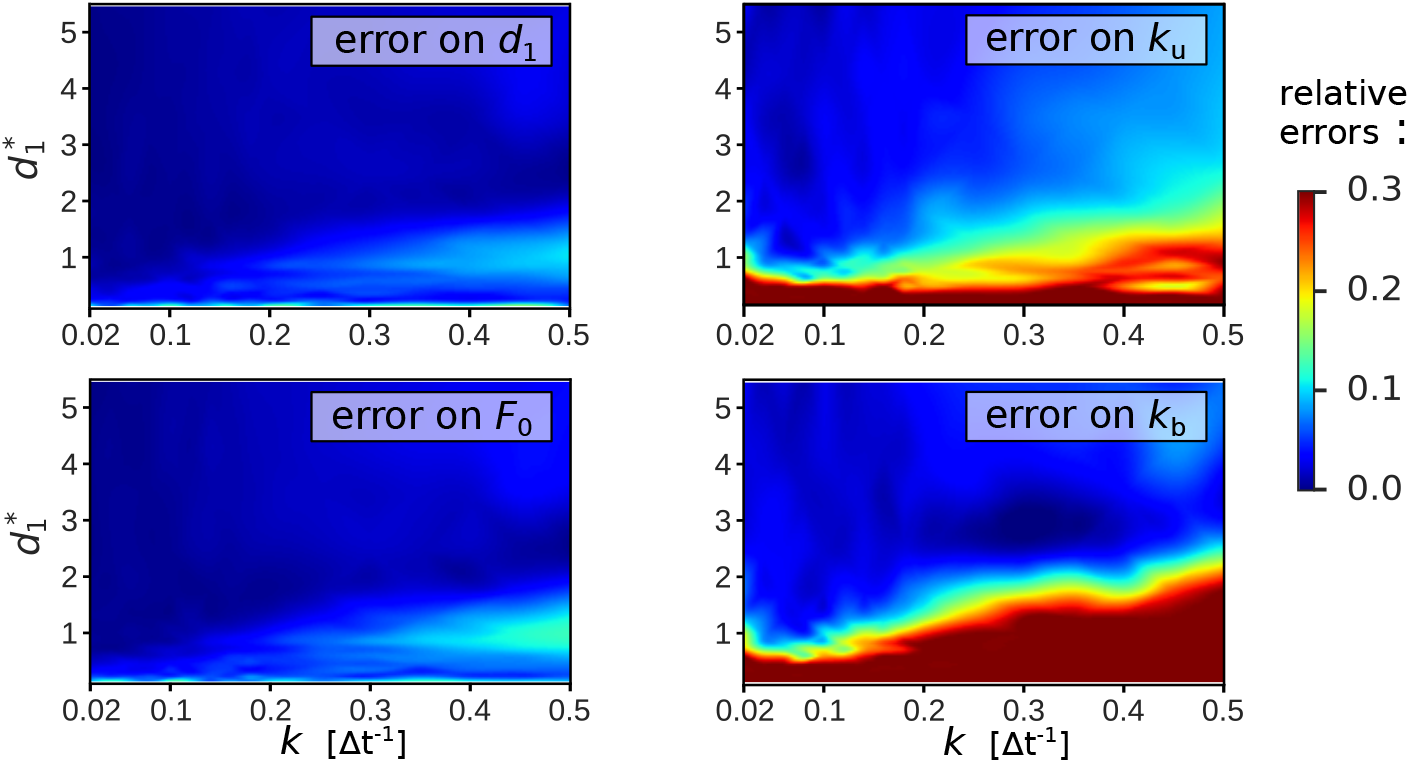
ExTrack parameter fits for a symmetric two-state model without sub-steps. Heat map of mean relative error on extracted parameters from the same simulations as in Fig. 1, but inferred with no sub-steps. ExTrack settings: no sub-steps, window length = 10.

**Supplementary Figure 3.**
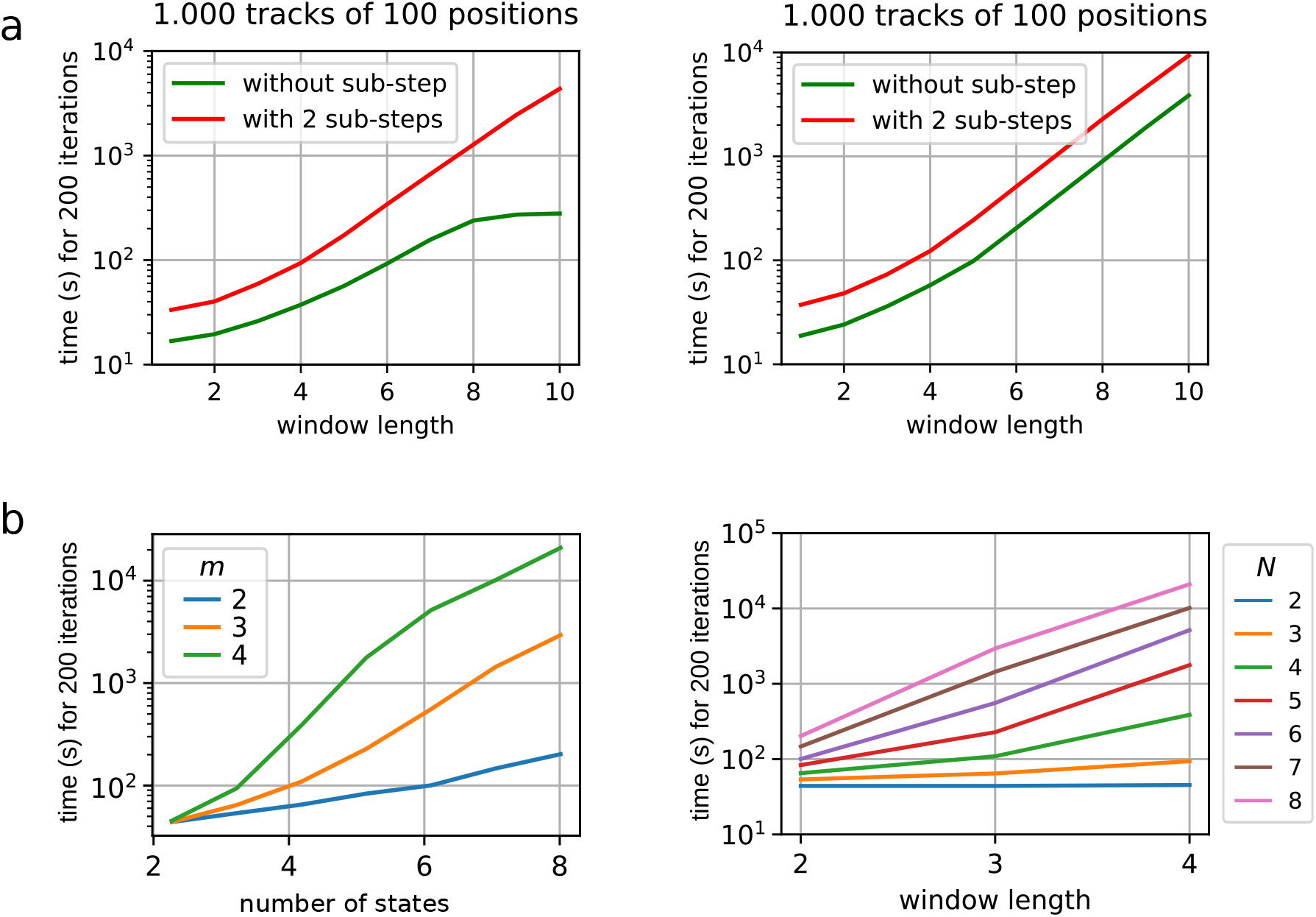
Computational time of ExTrack depending on sub-steps, window length and number of states. Computational time of ExTrack fitting module for 200 iterations (typical number of iterations needed for the fit of a two-state model) depending on the window length *m* from 1 to 10: Without sub-steps or with 2 sub-steps for tracks of 10 positions or 100 positions (**a**) or depending on the number of states *N* (**b**). with multiprocessing, 7 cores (see Methods section E).

**Supplementary Figure 4.**
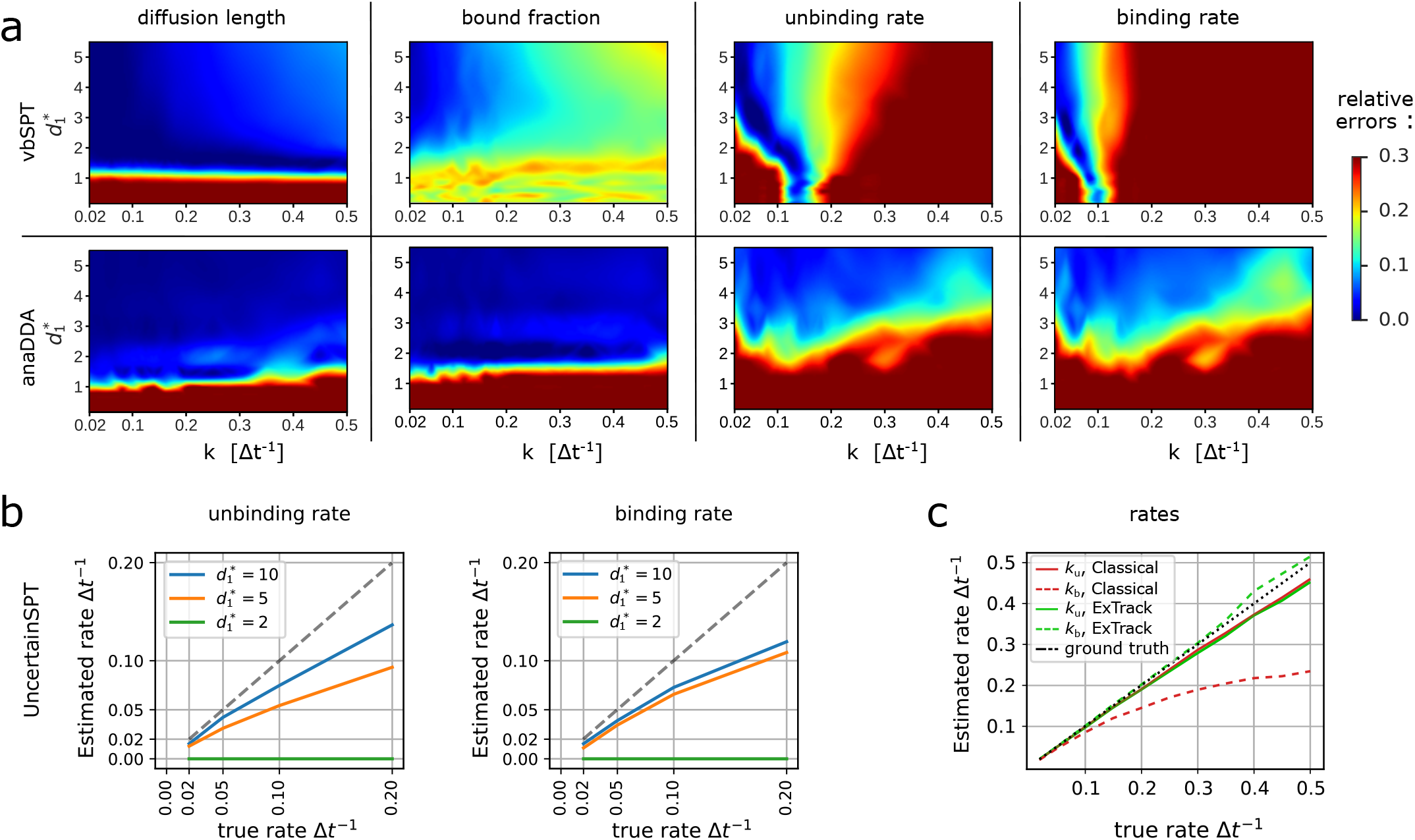
Error on two-state model parameters for different methods. **a**: Heat maps of mean relative error on extracted parameters from the same two-state simulations as in Fig. 1 for vbSPT and anaDDA. **b-c**: Plots of estimated transition rates as a function of the actual rates for a subset of the simulations in a. **b**: Results from UncertainSPT for different diffusion lengths. **c**: Results from a modified version of Extrack with a time-discretization approach (labeled Classical), which assumes transitions to occur at time points of molecule observations, and with our approach (labeled ExTrack), which assumes transitions to occur at the middle of each time steps (see Methods section A.2). Tracks simulated with 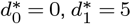.

**Supplementary Figure 5.**
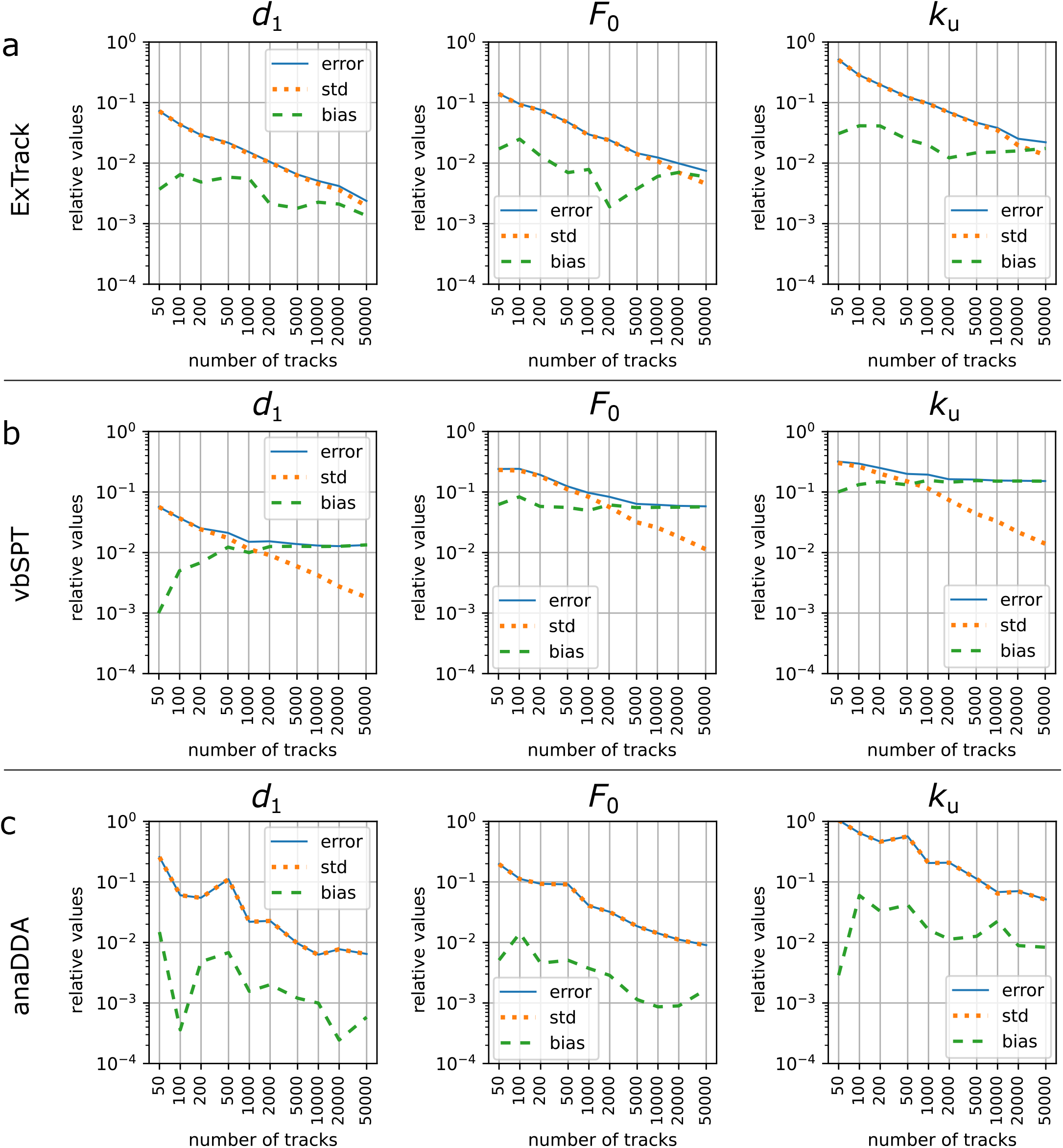
Identifying the sources of error of the different methods. Error, standard deviation and bias of parameters predicted by ExTrack (**a**), vbSPT (**b**) and anaDDA (**c**) depending on the number of tracks in case of two-state tracks (5 positions per track, *k*_u_ = *k*_b_ = 0.1 Δ*t*^*−*1^, 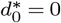 and 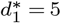). The error (RMSE) can be decomposed into bias (absolute value of the difference between the average estimate from all replicates and true parameter) and standard deviation (std) of the estimated parameters. 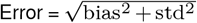. Obtained from 100 replicates. Here, all estimated values are relative to their true value. ExTrack settings: number of sub-steps = 2, window length = 10.

**Supplementary Figure 6.**
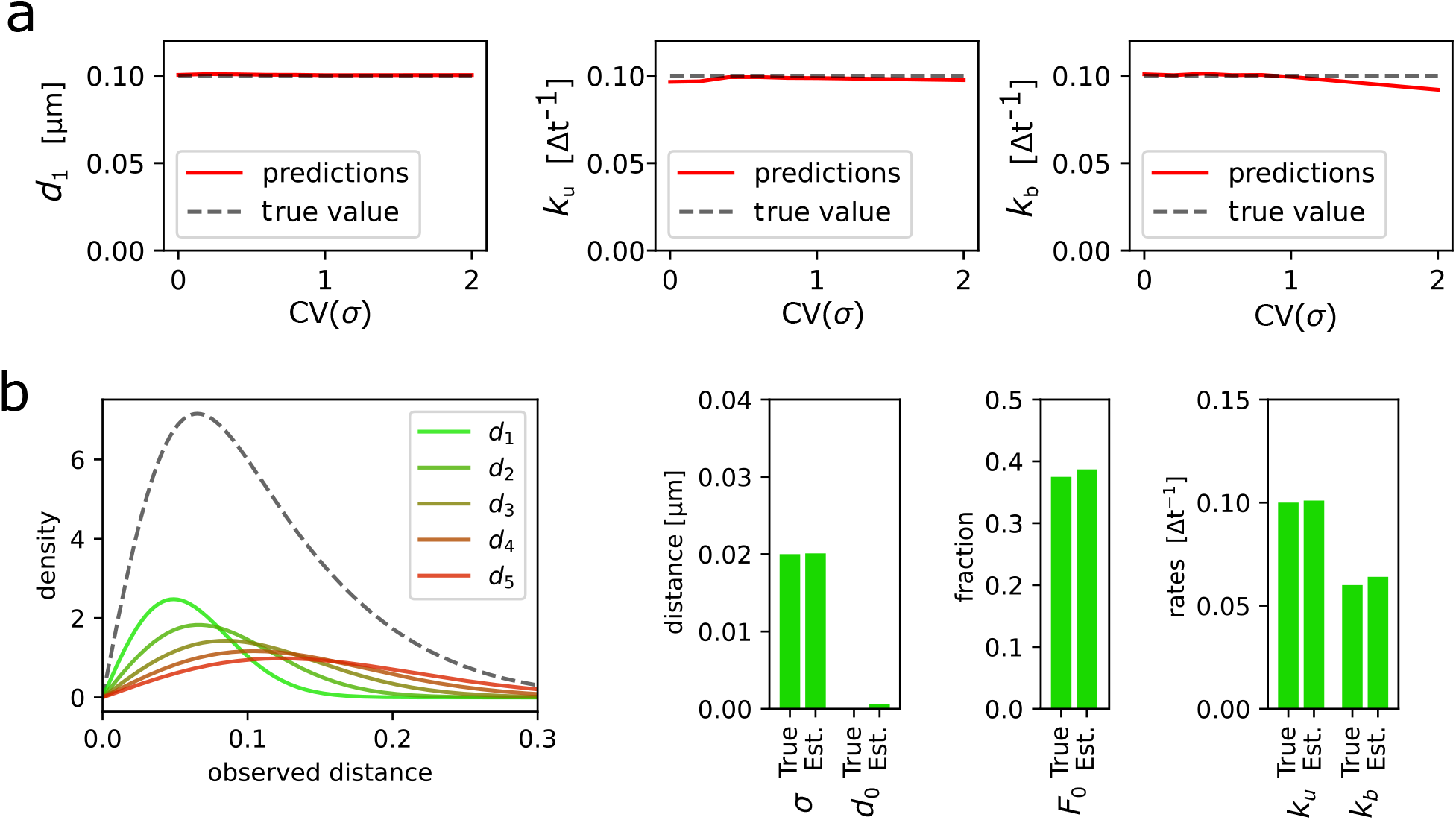
Robustness of ExTrack to biases due to distributions of diffusion coefficients or localization error. **a**: Predictions of *d*_1_, *k*_u_ and *k*_b_ in case of two-state parameter fits to two-state simulations with one immobile state and one diffusive state. Position at each time point show variable localization errors *σ*. Peak-wise localization errors were specified to the model. *σ* followed a chi-square distributions re-scaled so the mean localization error equals 0.02 µm (for sample distributions see inset of Fig. 2b). Simulations with *d*_1_ = 0.1*µ*m and *k* = *k*_u_ = *k*_b_ = 0.1. 10 replicates per condition. ExTrack settings: window length = 7, no sub-steps. **b**: We considered tracks from simulated particles with one immobile state 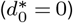 and five diffusive states with similar diffusion lengths of values 0.04, 0.06, 0.08, 0.1 and 0.12 µm (corresponding to *d*^*∗*^ from 2 to 12), all transition rates between each pair of diffusive states equal 0.1 Δ*t*^*−*1^ (resulting in average lifetimes of 2 Δ*t* for each diffusive state). This model results in indistinguishable diffusive tracks. Left: Distribution of displacements (for each dimension) of the five diffusive states. The grey dashed line represents the sum of the distributions. Right: Bar plots of true and estimated parameters obtained from fitting to a three-state model followed by aggregation of the diffusive states and computation of the resulting parameters. Here, the fractions are the global fractions computed from rates. See Table S2 for more details.

**Supplementary Figure 7.**
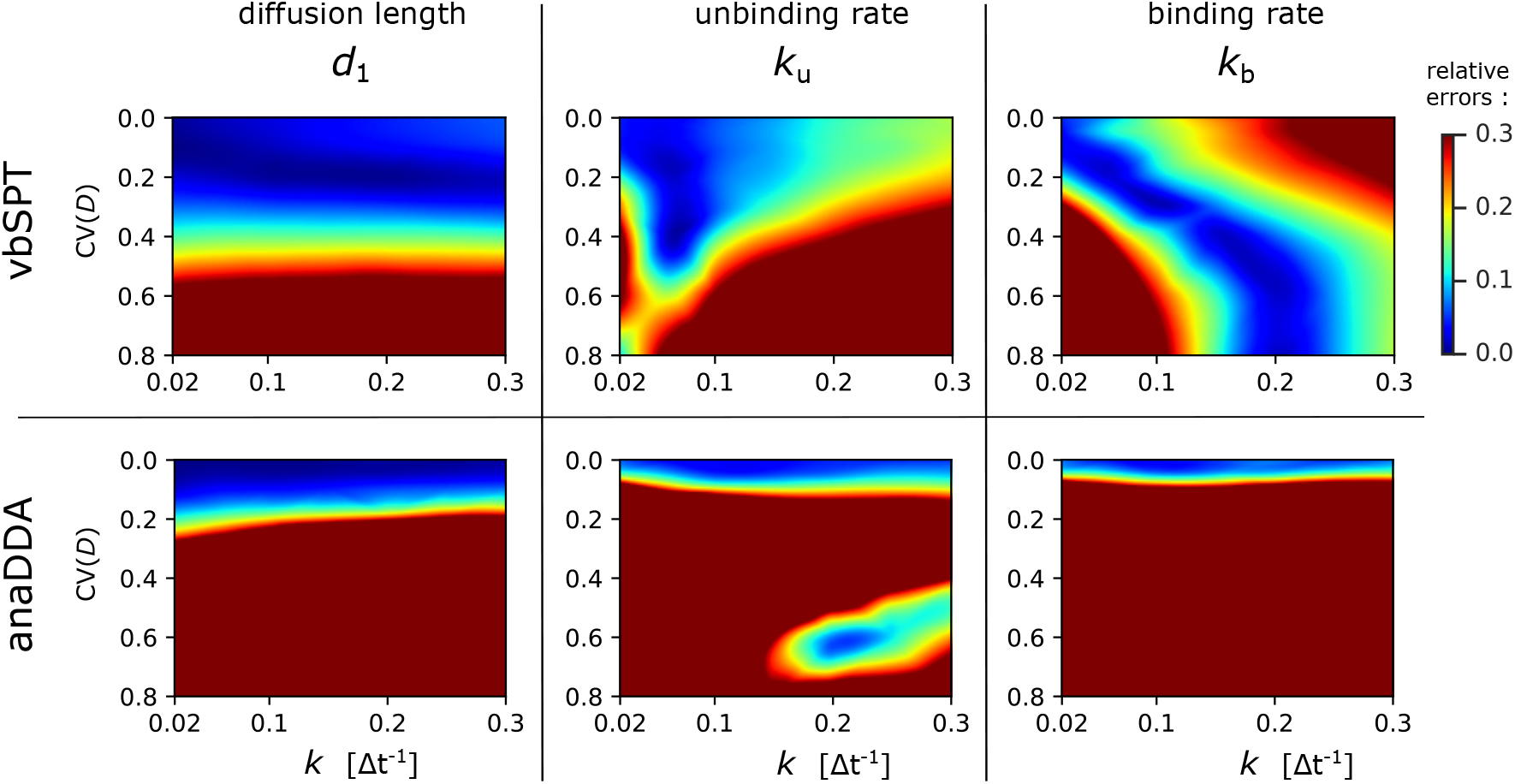
Robustness of vbSPT and anaDDA to biases due to a distribution of diffusion coefficients. Heatmaps of relative errors on *d*_1_, *k*_u_ and *k*_b_ with variable diffusion coefficient following the same protocol than in Fig. 2b for vbSPT and anaDDA.

**Supplementary Figure 8.**
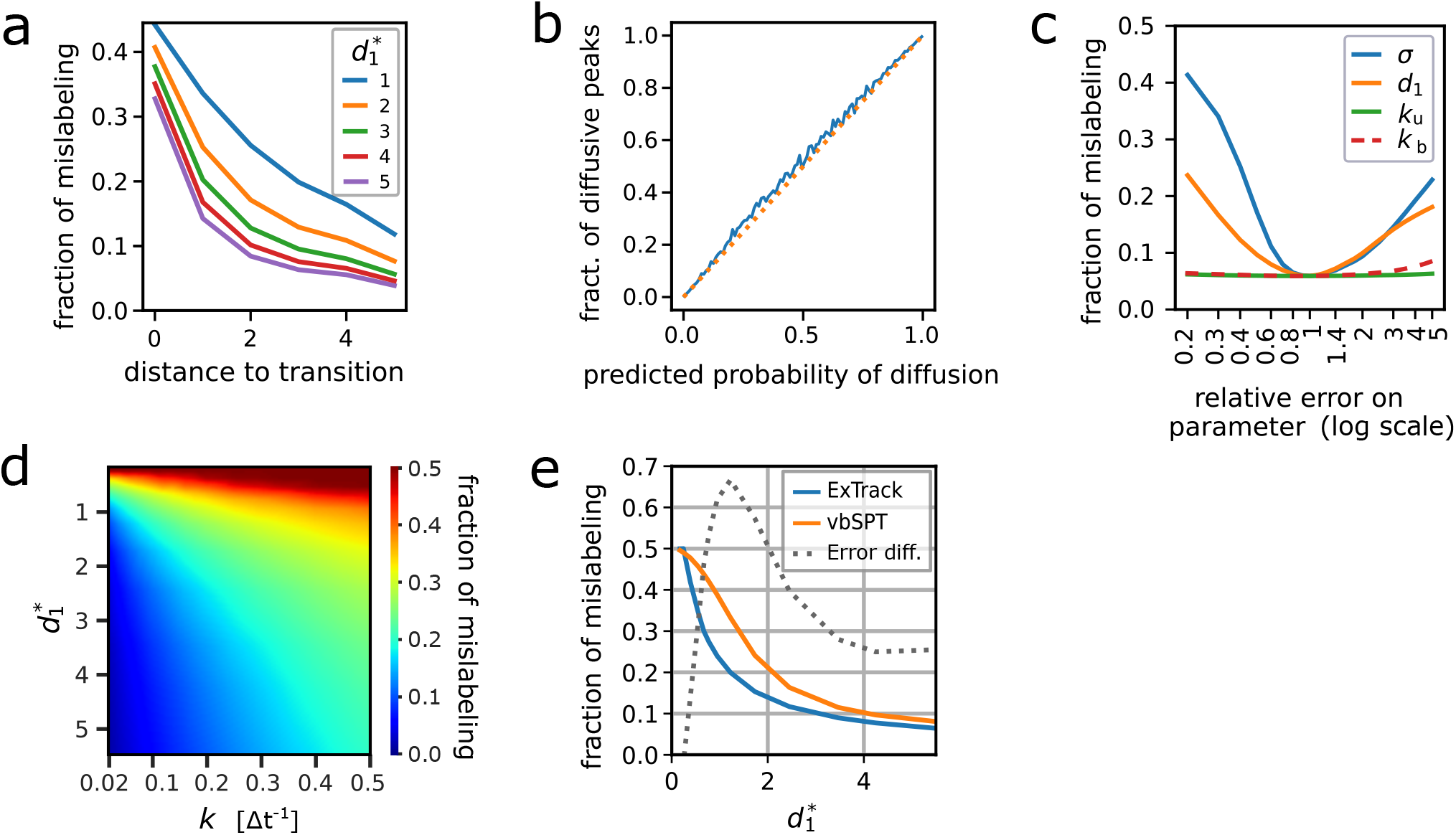
Capacity of the annotation module. Assessment of the annotation module accuracy by comparing state estimations (either probabilistic or categorical) with ground truth from simulated tracks. If not stated otherwise, 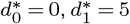 and *k*_u_ = *k*_b_ = 0.1 Δ*t*^*−*1^. **a**,**c-e**: Categorical state predictions are obtained by picking the most likely state for each time point. The fraction of mislabeled time points can thus be computed by comparison to the known true states. **a**: Fraction of mislabeled time points depending on (temporal) distance to transition time points. **b**: Fraction of time points actually in diffusive state depending on the probability to be diffusive estimated by ExTrack. More specifically, time points are binned according to their probability to be diffusive (*x*-axis) and for each bin we computed the fraction actually in diffusive state (*y*-axis). Binning of 0.01. **c**: Fractions of mislabeled time points using correct parameters except for one of them specified in the legend. X axis: relative error of the varied parameter compared to the true value underlying the simulated tracks. For this particular simulation, we used 10 000 tracks of 20 positions. Window length of 10. **d**: Heatmap of the fractions of mislabeled time points depending on 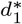 and *k*. **e**: Fraction of mislabeled time points depending on 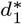 for ExTrack and vbSPT. The grey dotted curve (annotated as Error dif.) is the relative difference of the fraction of mislabeled time points between vbSPT and ExTrack (error _vbSPT_ - error _ExTrack_) / error _ExTrack_.

**Supplementary Figure 9.**
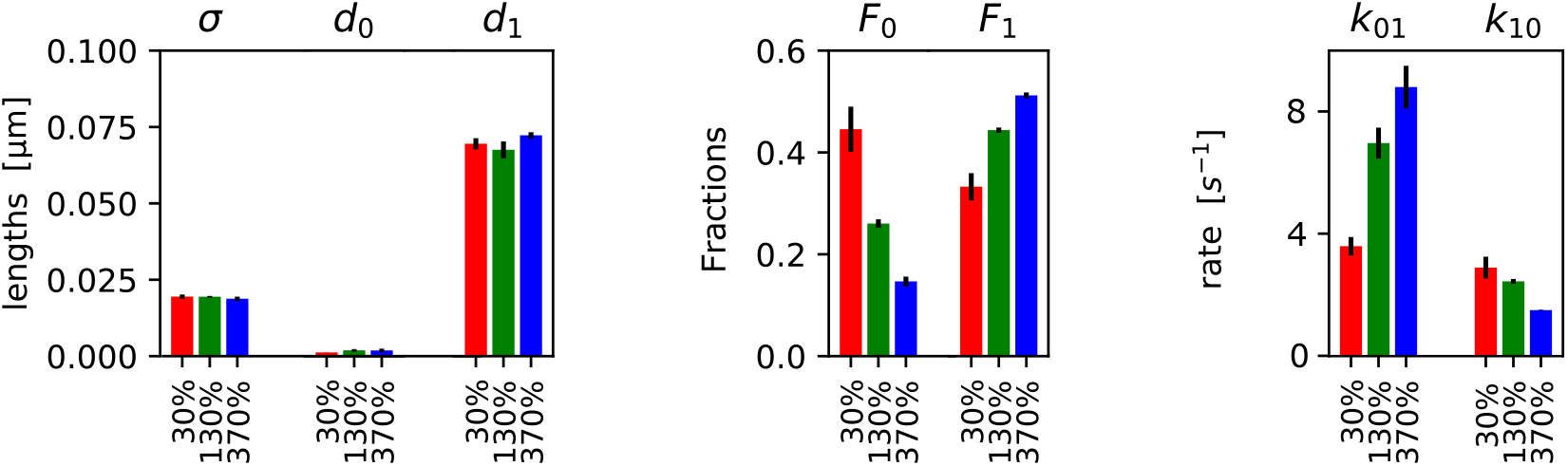
2-state model fits of PBP1b data. Results from parameter fits to experimental tracks of GFP-PBP1b of at least 3 time points (and considering not more than the first 50 time points assuming 2 states in ExTrack (3 replicates per condition, each replicate has at least 17.000 tracks of average lifetime from 6.3 to 7.5 positions). ExTrack settings: Window length = 4, no sub-steps. State fractions obtained from rates. Error bars: standard deviations between replicates.

**Supplementary Figure 10.**
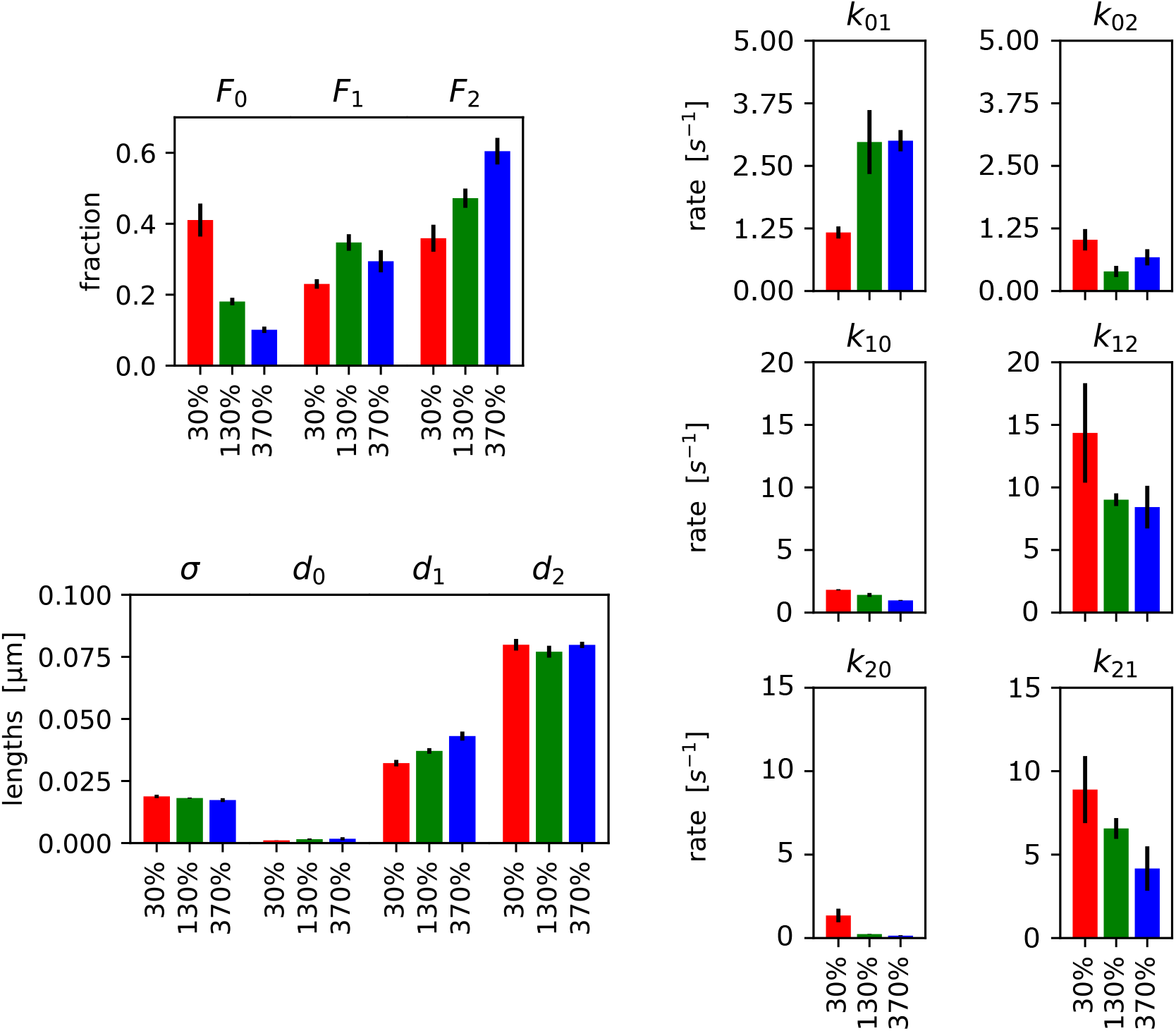
3-state model fits of PBP1b data. Results from parameter fits to the same experimental tracks of GFP-PBP1b considered in Fig. 9 but assuming 3 states in ExTrack. ExTrack settings: Window length = 4. State fractions obtained from rates. Error bars: standard deviations between replicates.

**Supplementary Figure 11.**
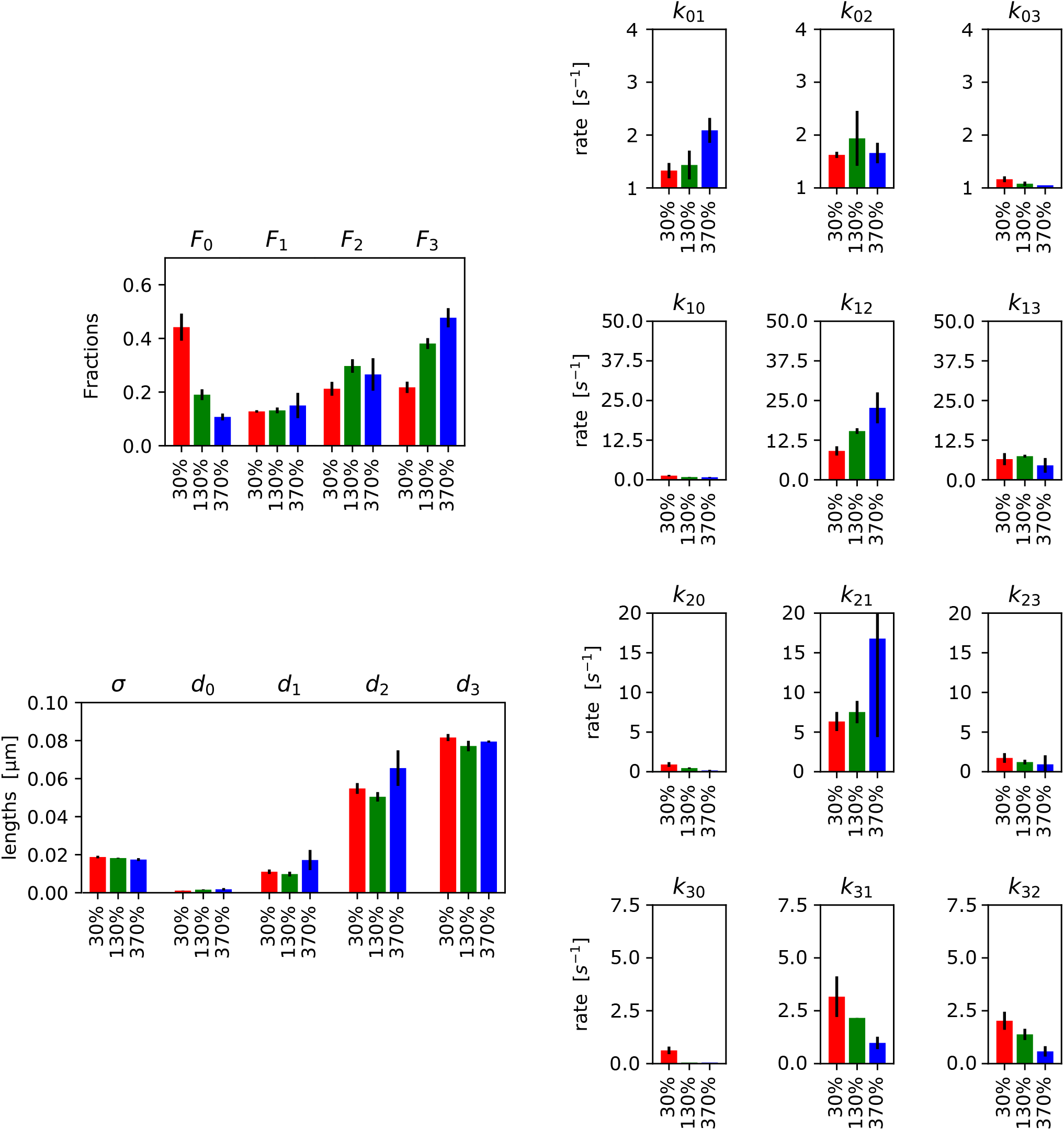
4-state model fits of PBP1b data. Results from parameter fits to the same experimental tracks of GFP-PBP1b considered in Fig. 9 but assuming 4 states in ExTrack. Window length = 5. State fractions obtained from rates. Error bars: standard deviations between replicates.

**Supplementary Figure 12.**
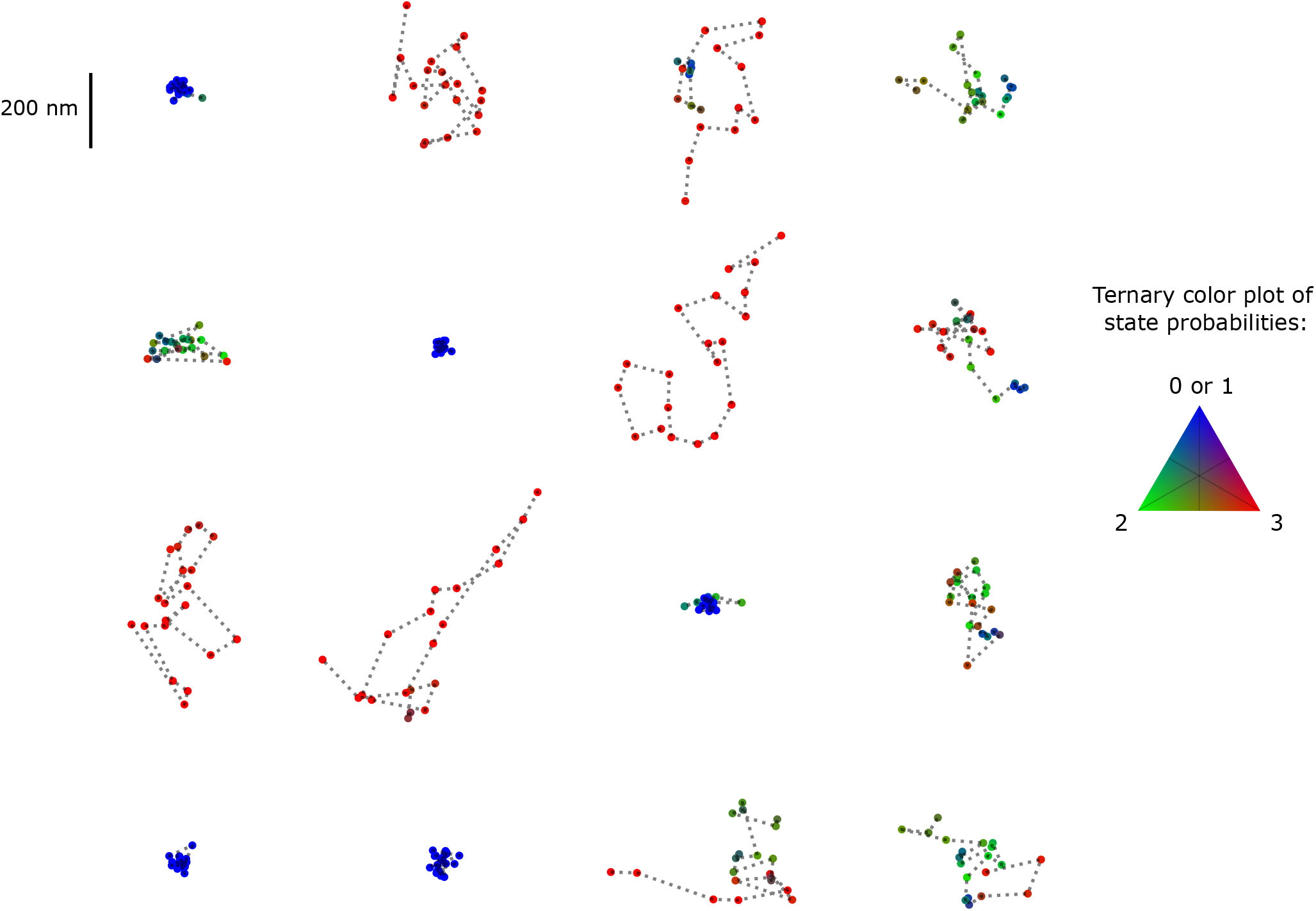
State annotations of GFP-PBP1b using a 4-state model. State annotation from parameters obtained from the 4-state fits to GFP-PBP1b data at 130% PBP1b expression. Immobile (state 0 and 1) in blue, intermediate diffusion state (state 2) in green and diffusive state (state 3) in red. The ternary plot represents the intermediate probabilities. ExTrack settings: window length = 7, no sub-steps. Annotation using parameter obtain from ExTrack fits on the same data. *d*_1_ is so low that on the short timescale of transient binding molecules are nearly immobile.

**Supplementary Figure 13.**
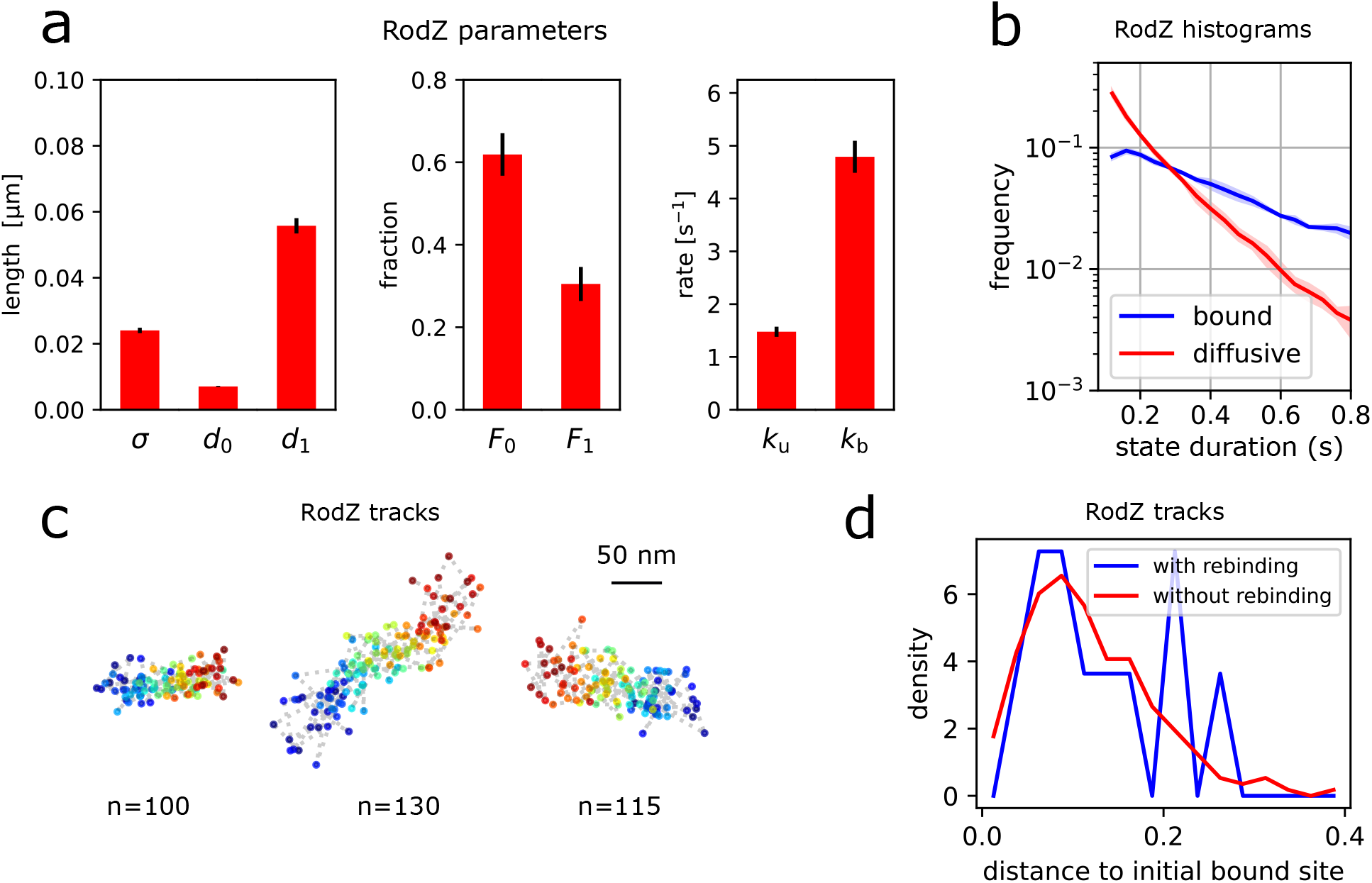
Complementary results for GFP-RodZ and GFP-PBP1b tracks. **a-c**: ExTrack analysis of GFP-RodZ data (same as in Fig. 4f-h) **a**: Parameters found by ExTrack for GFP-RodZ tracks (3 replicates, each replicate has at least 25.000 tracks of average lifetime from 6.3 to 7.4 positions), considering all tracks with at least 3 time points and restricting analysis to the 50 first time points. **b**: Histogram of the time spent in bound or diffusive states, among tracks of at least 21 time points using parameters from **a. c**: Examples of long bound RodZ tracks in linear motion. **d**: PBP1b rebinding. Histograms of the distances in between initially bound position and after 4 diffusive steps of PBP1b tracks that are subsequently either rebinding (blue) or not rebinding (red) (see Methods). Histograms show no noticable difference. Error bars and shaded regions: standard deviations between replicates.

**Supplementary Figure 14.**
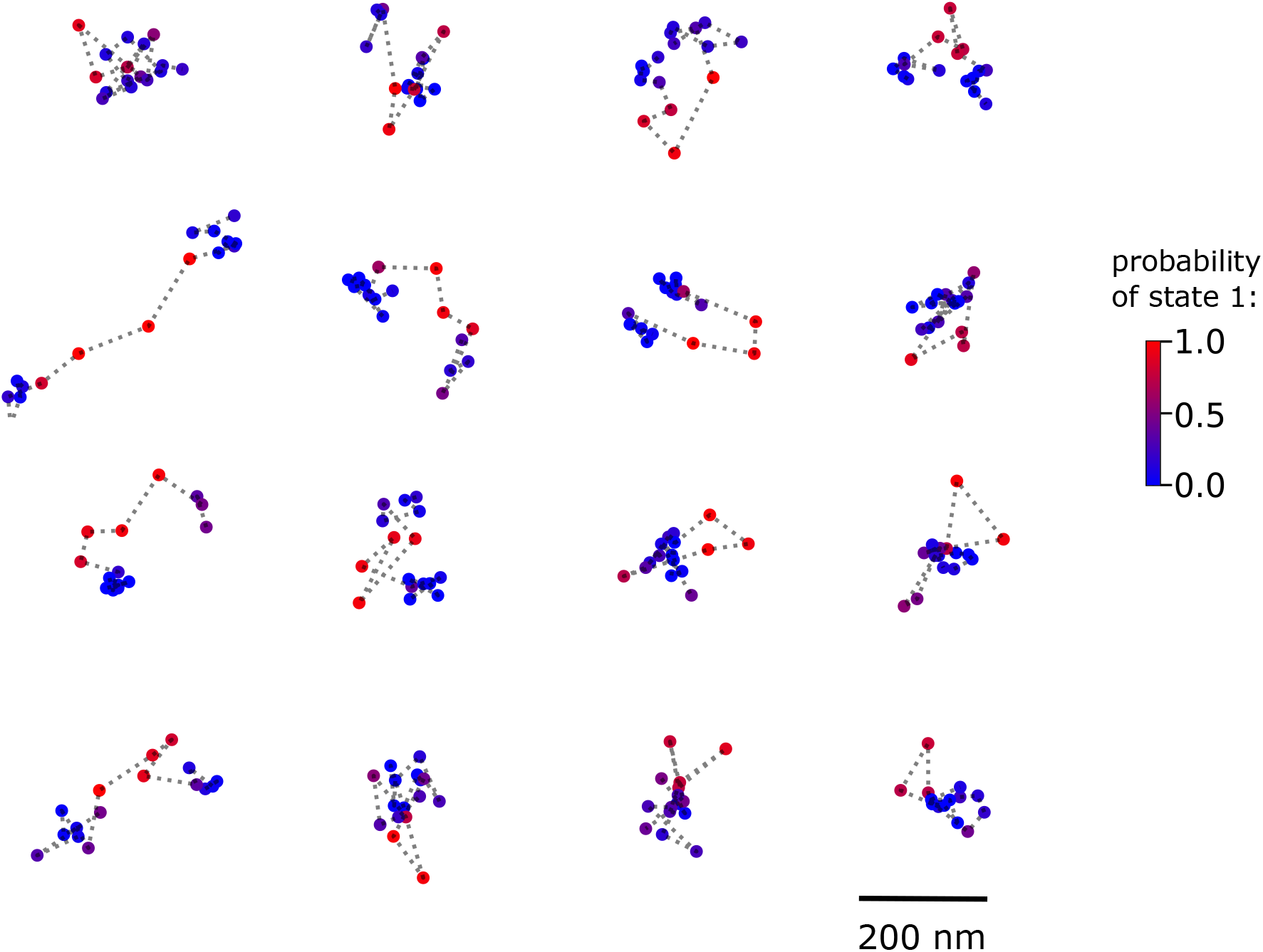
State annotation or RodZ tracks that show rebinding. Two-state annotations of GFP-RodZ initially immobile (at least 3 positions), then diffusive for 4 positions and then immobile again. Annotation using parameters obtained from fits on the same data. Immobile (state 0) in blue and diffusive (state 1) in red. ExTrack settings: window length = 10. Colorbar: probability of state 1 (diffusive).

## Supplemental Tables

**Supplementary Table 1.** Here, we use ExTrack to fit parameters of a three-state model to different simulations of three-state data with qualitatively different types of transitions. Model 1: immobile state, intermediate diffusion state and high diffusion state. 10 000 tracks of 10 positions. Model 2 to 4 represent harder models with two immobile states and one diffusive state (10 000 tracks of 50 positions each). They all have a transient immobile state (state 0), a stable immobile state (state 1) and a diffusive state (state 2). Model 2: transitions from state 0 to 1 to 2 to 0 in a circular fashion. Model 3: transitions from state 2 to 1 to 0 to 2 in a circular fashion. Model 4: more complex transitions between states.

| 3-state models | | $\sigma$ | $d_0$ | $d_1$ | $d_2$ | $F_0$ | $F_1$ | $F_2$ | $k_{01}$ | $k_{02}$ | $k_{10}$ | $k_{12}$ | $k_{20}$ | $k_{21}$ |
| --- | --- | --- | --- | --- | --- | --- | --- | --- | --- | --- | --- | --- | --- | --- |
| 1 | True | 0.020 | 0.000 | 0.040 | 0.100 | 0.333 | 0.333 | 0.333 | 0.100 | 0.100 | 0.100 | 0.100 | 0.100 | 0.100 |
|  | Estimated | 0.020 | 0.001 | 0.040 | 0.101 | 0.342 | 0.330 | 0.328 | 0.097 | 0.093 | 0.098 | 0.093 | 0.098 | 0.098 |
| 2 | True | 0.020 | 0.000 | 0.000 | 0.140 | 0.069 | 0.517 | 0.414 | 0.300 | 0.000 | 0.000 | 0.040 | 0.050 | 0.000 |
|  | Estimated | 0.020 | 0.000 | 0.000 | 0.141 | 0.069 | 0.518 | 0.413 | 0.297 | 0.005 | 0.000 | 0.040 | 0.050 | 0.000 |
| 3 | True | 0.020 | 0.000 | 0.000 | 0.140 | 0.077 | 0.462 | 0.462 | 0.000 | 0.300 | 0.050 | 0.000 | 0.000 | 0.050 |
|  | Estimated | 0.020 | 0.000 | 0.000 | 0.141 | 0.089 | 0.494 | 0.417 | 0.075 | 0.190 | 0.047 | 0.009 | 0.000 | 0.051 |
| 4 | True | 0.020 | 0.000 | 0.000 | 0.140 | 0.060 | 0.489 | 0.451 | 0.250 | 0.050 | 0.020 | 0.040 | 0.040 | 0.010 |
|  | Estimated | 0.020 | 0.000 | 0.000 | 0.141 | 0.046 | 0.504 | 0.450 | 0.358 | 0.046 | 0.000 | 0.041 | 0.042 | 0.009 |

**Supplementary Table 2.** 2-state and 3-state fits of tracks from simulated particles either in immobile state (state 0) or in one of 5 diffusive states (states 1 to 5). Here, unbinding rates *k*_0,*j*_ = 0.02 Δ*t*^*−*1^, binding rates *k*_0,*j*_ = 0.06 Δ*t*^*−*1^ and other rates *k*_*i,j*_ = 0, 0.02 or 0.1 Δ*t*^*−*1^ for *i* and *j* from 1 to 5. *, *k*_u_ and *k*_b_ are the global unbinding and binding rates, respectively, obtained as the sum of the unbinding rates, and the average of the binding rates weighted by the fractions in diffusive state. ExTrack settings: window length = 6. All distances in µm and rates in Δ*t*^*−*1^.

| ExTrack Model | Set of $d$ | $k_{i,j}$ | Parameters | $\sigma$ | $d_0$ | $d_1$ | $F_0$ | $k_u^*$ | $k_b^*$ |
| --- | --- | --- | --- | --- | --- | --- | --- | --- | --- |
| 2 states | (0.04, 0.06, 0.08, 0.10, 0.12) | 0.10 | Estimated | 0.021 | 0.001 | 0.090 | 0.458 | 0.130 | 0.110 |
|  |  |  | True | 0.020 | 0.000 | - | 0.375 | 0.1 | 0.06 |

**Supplementary Table 2.** 2-state and 3-state fits of tracks from simulated particles either in immobile state (state 0) or in one of 5 diffusive states (states 1 to 5).
| ExTrack Model | Set of $d$ | $k_{i,j}$ | Parameters | $\sigma$ | $d_0$ | $d_1$ | $d_2$ | $F_0$ | $k_u$ | $k_b$ |
| --- | --- | --- | --- | --- | --- | --- | --- | --- | --- | --- |
| 3 states | (0.04, 0.06, 0.08, 0.10, 0.12) | 0.00 | Estimated | 0.020 | 0.001 | 0.054 | 0.107 | 0.395 | 0.101 | 0.066 |
|  |  | 0.02 | Estimated | 0.020 | 0.001 | 0.053 | 0.107 | 0.387 | 0.101 | 0.064 |
|  |  | 0.10 | Estimated | 0.020 | 0.000 | 0.053 | 0.110 | 0.399 | 0.102 | 0.068 |
|  | (0.04, 0.08, 0.12, 0.16, 0.20) | 0.02 | Estimated | 0.020 | 0.000 | 0.064 | 0.174 | 0.405 | 0.105 | 0.072 |
|  |  |  | True | 0.020 | 0.000 | - | - | 0.375 | 0.100 | 0.060 |

